# Derlin Dfm1 Employs a Chaperone Function to Resolve Misfolded Membrane Protein Stress

**DOI:** 10.1101/2022.01.25.477788

**Authors:** Rachel Kandel, Jasmine Jung, Della Syau, Tiffany Kuo, Livia Songster, Analine Aguayo, Sascha Duttke, Christopher Benner, Sonya Neal

## Abstract

Accumulation of misfolded proteins is a known source of cellular stress and can be detrimental to cellular health. While protein aggregation is a known hallmark of many diseases, the mechanisms by which protein aggregates cause toxicity and the molecular machines that prevent this toxicity are not completely understood. Here, we show that the accumulated misfolded membrane proteins form endoplasmic reticulum (ER) localized aggregates, impacting ubiquitin and proteasome homeostasis. Additionally, we have identified a chaperone ability of the yeast rhomboid pseudoprotease Dfm1 to influence solubilization of misfolded membrane proteins and prevent toxicity from misfolded membrane proteins. We establish that this function of Dfm1 does not require recruitment of the ATPase Cdc48 and it is distinct from Dfm1’s previously identified function in dislocating misfolded membrane proteins to the cytosol for degradation.

## INTRODUCTION

Proper and efficient protein folding is essential for maintaining cellular health. Eukaryotic cells are equipped with protein quality control pathways for preventing the accumulation of misfolded proteins. One of the major pathways of protein quality control at the endoplasmic reticulum (ER) is ER associated degradation (ERAD)^1^. ERAD utilizes the ubiquitin proteasome system (UPS) to selectively target and degrade unassembled or misfolded proteins at the ER^2^. To properly regulate the large variety of proteins folded at the ER, there are multiple branches of ERAD, with specific machinery that is exclusive to each branch. ERAD-L targets misfolded luminal proteins while ERAD-M and ERAD-C target misfolded membrane proteins^3, 4^.

ERAD is a well conserved process from yeast to mammals. ERAD of membrane proteins requires four universal steps: 1) substrate recognition^5^, 2) substrate ubiquitination^6^, 3) retrotranslocation of substrate from the ER to the cytosol^7^, and 4) degradation by the cytosolic proteasome^2^. A hexameric cytosolic ATPase, Cdc48 in yeast and p97 in mammals, is required for retrotranslocation of all ERAD substrates^8,9,10^. While similar machinery is needed for all branches of ERAD, there are some distinct differences. ERAD-M substrates can be targeted by the DOA (degradation of alpha2) pathway or the HRD pathway (Hydroxymethyl glutaryl-coenzyme A reductase degradation), utilizing the E3 ligases Doa10 and Hrd1, respectively. Dfm1, a yeast rhomboid pseudoprotease, is specifically required for the retrotranslocation of misfolded membrane substrates, in both the HRD and DOA pathways^7^. The HRD pathway is also required for degradation of ERAD-L substrates. The other yeast rhomboid pseudoprotease, Der1, is required for retrotranslocation of luminal proteins in yeast^11^. The retrotranslocation function of Der1 requires the E3 ligase Hrd1, with each protein forming half of a channel for misfolded protein dislocation^12^.

In contrast to Der1, Dfm1’s retrotranslocation function is independent of Hrd1^7, 13^. We have previously observed that in *dfm1Δ* cells, when a misfolded membrane protein is strongly expressed, the cells show a severe growth defect^13^. This is seen specifically in the absence of Dfm1, and this growth defect is not observed in the absence of other ERAD components, indicating a specific function for Dfm1 in sensing and/or adapting cells to misfolded membrane protein stress (Fig. 1)^13^. This is in line with a previous study linking Dfm1 to ER homeostasis^14^.

**Figure 1:**
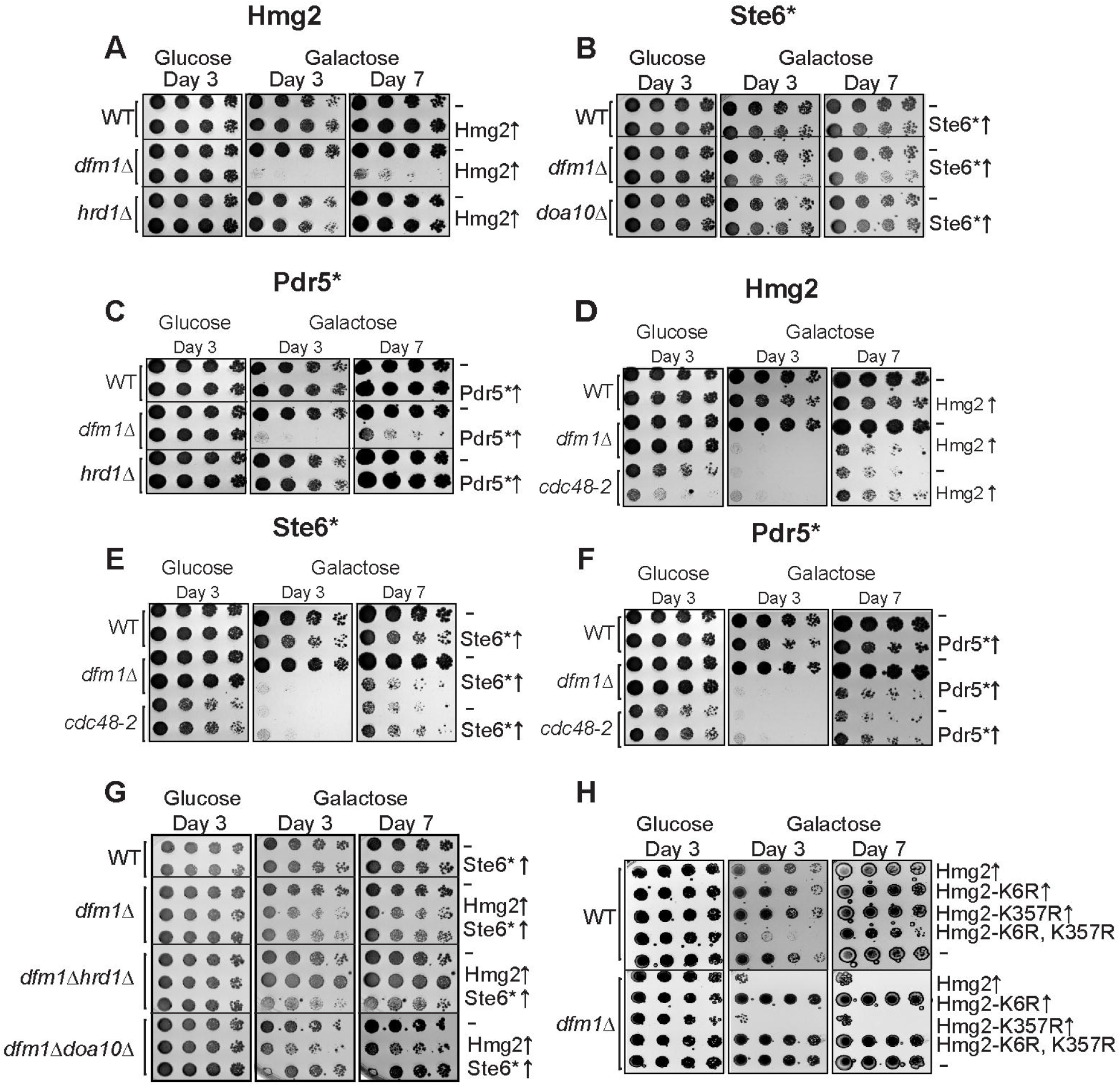
**Integral Membrane Protein Overexpression Causes a Growth Defect in *dfm1Δ* Cells in an ERAD Independent Manner** (A) WT, *dfm1*Δ, and *hrd1*Δ cells containing either GAL_pr_-HMG2-GFP or EV were compared for growth by dilution assay. Each strain was spotted 5-fold dilutions on glucose or galactose-containing plates to drive HMG2-GFP overexpression, and plates were incubated at 30°C. (B) Dilution assay as described in (A) except using WT, *dfm1*Δ, and *doa10*Δ cells containing either GAL_pr_-STE6*-GFP or EV. (C) Dilution assay as described in (A) except using WT, *dfm1*Δ, and *hrd1*Δ cells containing either GAL_pr_-PDR5*-HA or EV. (D) Dilution assay as described in (A) except using WT *dfm1*Δ, and *cdc48-2* cells. (E) Dilution assay as described in (B) except using WT *dfm1*Δ, and *cdc48-2* cells. (F) Dilution assay as described in (C) except using WT *dfm1*Δ, and *cdc48-2* cells. (G) Dilution assay as described in (A) except using WT, *dfm1*Δ, *dfm1*Δ*hrd1*Δ and *dfm1*Δ*doa10*Δ cells containing either GAL_pr_-Hmg2-GFP, GAL_pr_-STE6*-GFP, or EV. (H) Dilution assay as described in (A) except using WT and *dfm1*Δ cells containing either GAL_pr_-Hmg2-GFP, GAL_pr_-Hmg2 (K6R)-GFP, GAL_pr_-Hmg2 (K357R)-GFP, GAL_pr_-Hmg2 (K6R and K357R)-GFP or EV.

While misfolded proteins are known to pose a threat to the health of cells, in many circumstances, the exact mechanism by which these proteins prove toxic to cells is unclear. Cellular stress responses to misfolded proteins have been studied extensively in the context of misfolded luminal ER proteins, in the form of the unfolded protein response (UPR). In contrast, there is very little research into how accumulation of misfolded membrane proteins both affects cells and how cells prevent toxicity from these proteins. Despite the dearth of research on this topic, a previous study implicates the transcription factor Rpn4^15^ in resolving misfolded membrane protein stress, a protein we have also identified in the present study. In addition, we show that rhomboid pseudoprotease Dfm1 as well as the deubiquitinases Ubp6, Doa4, Ubp14, and Ubp9 are critical in preventing misfolded membrane protein toxicity. For Dfm1, we determine that its ability to prevent membrane protein toxicity is because of a previously unidentified chaperone function. This study is the first to demonstrate any rhomboid protein acting as a chaperone. Furthermore, our results indicate that Dfm1 does not have to recruit the ATPase Cdc48 to function as a chaperone. We propose a model in which upon accumulation of misfolded membrane proteins in the absence of Dfm1, misfolded membrane proteins form aggregates, resulting in disruptions to ubiquitin homeostasis and impairment to proteasomes. In the presence of Dfm1, this toxicity is prevented by Dfm1’s ability to solubilize membrane proteins, independent of its ability to retrotranslocate proteins.

## RESULTS

### Absence of Dfm1 and Expression of Integral Misfolded Membrane Proteins Cause Growth Stress

Previous research from the Neal lab has revealed that accumulation of a misfolded membrane protein in the absence of Dfm1 causes a severe growth defect in the substrate-toxicity assay^16^. In the substrate-toxicity assay, strains with a misfolded protein under the control of a galactose inducible promoter are plated in a spot assay onto selection plates with either 2% galactose or 2% dextrose as a carbon source (Fig. 1)^17^. This allows for comparison of growth of yeast strains with different genetic perturbations with expression of misfolded substrates. This growth defect can be seen with both strong expression of three misfolded membrane proteins; Hmg2 and Pdr5*, ERAD-M substrates, as well as Ste6*, an ERAD-C substrate (Fig. 1A-C). We have previously shown that this growth defect is specific to misfolded membrane proteins at the ER, as expression of an ERAD-L substrate, CPY*, elicits no growth defect and still shows the same growth as WT cells with CPY*^16^. Interestingly, this growth defect is not seen when misfolded proteins are expressed in the absence of other ERAD components, such as the E3 ligases Hrd1 and Doa10 (Fig.1A-C). In the case of *dfm1Δ*, *hrd1Δ,* and *doa10Δ* cells, misfolded membrane proteins accumulate at the ER due to defects in ERAD, but only in the case of *dfm1Δ* cells is a growth defect observed with Hmg2 expression. Altogether, we surmise that this growth defect triggered by the absence of Dfm1 along with expression of misfolded membrane protein is due to cellular stress caused by misfolded membrane protein toxicity.

By utilizing the substrate-toxicity assay, we observed a growth defect in *dfm1Δ* cells and normal growth in *hrd1Δ* and *doa10Δ* cells upon expression of ERAD membrane substrates. The cell biological difference amongst these ERAD knockout strains is that membrane substrates are ubiquitinated in *dfm1Δ* cells and not ubiquitinated in *hrd1Δ* and *doa10Δ* cells, due to the absence of the ER E3 ligases. One possibility is that the growth stress is not specific to *dfm1Δ* cells and is solely dependent on the accumulation of ubiquitinated membrane substrates. To rule out this possibility, we utilized a hypomorphic Cdc48 allele, *cdc48-2*, which also results in the accumulation of ubiquitinated ERAD membrane substrates, just like *dfm1Δ* cells. The substrate-toxicity assay was employed on *cdc48-2* strains using membrane substrates Hmg2, Pdr5* (another ERAD-M substrate), and Ste6* (Fig. 1D-F). These strains showed a growth defect while growing on galactose plates due to inherent slow growth of *cdc48-2* strains, but this was not worsened by expression of misfolded integral membrane proteins, despite those membrane proteins being ubiquitinated. These results indicate that Dfm1 plays a specific role in the alleviation of misfolded membrane protein stress.

### Growth Defect in dfm1Δ Cells is Ubiquitination Dependent

The observation that a growth defect is only seen in the absence of Dfm1, and not in cells lacking either of the ER E3 ligases Hrd1 and Doa10, led us to hypothesize that this growth defect is dependent upon ubiquitination of the misfolded membrane proteins. The substrate-toxicity assay results using *cdc48-2* cells indicate that the growth defect is not solely due to defective ERAD or the accumulation of ubiquitinated misfolded membrane proteins. Nonetheless, we still explored the possibility that misfolded membrane protein-induced toxicity is dependent on substrate ubiquitination.

We examined whether growth defects were seen in either *dfm1Δhrd1Δ* or *dfm1Δdoa10Δ* cells expressing either Hmg2 (a Hrd1 target) or Ste6* (a Doa10 target), respectively (Fig. 1G). These results showed no growth defect in the double mutants for which the membrane protein expressed was not ubiquitinated by the absent E3 ligase: *dfm1Δhrd1Δ* cells expressing Hmg2 and *dfm1Δdoa10Δ* cells expressing Ste6* (Fig. 1G). In contrast, a growth defect was observed in the double mutants for which the absent E3 ligase did not participate in ubiquitination of the expressed membrane protein: *dfm1Δhrd1Δ* expressing Ste6* and *dfm1Δdoa10Δ* expressing Hmg2. This indicates that growth stress in *dfm1Δ* cells is dependent upon ubiquitination of the accumulated misfolded membrane protein.

As an alternative approach to determine if membrane proteins must be ubiquitinated to cause toxicity in the absence of Dfm1, we tested the expression of well-characterized, stabilized Hmg2 mutants. These mutants, Hmg2 (K6R), Hmg2 (K357R), and Hmg2 (K6R, K357R), were previously identified by the Hampton lab in a genetic screen for stabilized Hmg2 mutants^18^. Both KàR stabilized mutations disrupt Hmg2 ubiquitination, and these sites are hypothesized to be Hmg2 ubiquitination sites. While the Hampton lab has shown that ubiquitination levels of both substrates are negligible, they also showed that the K6R mutant is not further stabilized in an ERAD deficient background, while the K357R mutant is slightly more stable in an ERAD deficient background than in a WT background^18^. We propose that because of this slight level of degradation in the K357R mutant, some fraction of this mutant must be ubiquitinated and targeted to the Hrd1 ERAD pathway. Our model predicts that growth defect is ubiquitin dependent: thus, we would expect that more stabilized K6R with negligible ubiquitination should not elicit a growth defect whereas K357R, which slightly undergoes ubiquitination and degradation should elicit a growth defect. Indeed, we observed no growth defect in *dfm1Δ* expressing the K6R mutant, while the K357R mutant still showed a growth defect. Moreover, the growth defect is still observed in the double mutant Hmg2 (K6R, K357R), consistent with the model that growth stress in absent of Dfm1 is dependent on the accumulation of ubiquitinated membrane substrates (Fig. 1H).

### Growth Defect in dfm1Δ Cells is Not Caused by Activation of the Unfolded Protein Response

The canonical ER stress pathway triggered by the accumulation of misfolded proteins is the unfolded protein response (UPR)^19^. The UPR is known to be induced by the accumulation of misfolded soluble proteins within the ER lumen. To test if misfolded membrane protein accumulation at the ER activates UPR, we used a fluorescence-based flow cytometry assay. In this assay, yeast cells encoding both a galactose inducible misfolded protein and UPR optical reporter 4xUPRE-GFP were treated with or without 0.2% galactose and 2 µg/mL of the ER stress inducing drug tunicamycin or DMSO as the vehicle control. GFP expression was measured by flow cytometry every hour for 5 hours following galactose treatment. We found that GFP expression did not increase over the time course in *dfm1Δ* cells compared to *pdr5Δ* cells expressing any of the substrates tested: Hmg2 (ERAD-M), Ste6* (ERAD-C), or empty vector (EV) (Fig. 2A-H). We also determined that the stress in *dfm1Δ* cells is not due to a lack of ability of these cells to mount UPR, as addition of tunicamycin to these cells allowed them to activate UPR at similar levels as *pdr5Δ* cells (Fig. 2A-H). As expected, expression of the ERAD-L substrate CPY* activated the UPR in *dfm1Δ* cells and *pdr5Δ* cells (Fig. 2E&F). Unexpectedly, the level of UPR activation was much higher in *dfm1Δ* cells expressing CPY* than in *pdr5Δ* cells. This was further seen in the cells that were treated with tunicamycin, as CPY* expression with tunicamycin treatment in *dfm1Δ* cells resulted in a large percentage of the cells dying by the 5-hour time point, resulting in a decrease in average fluorescence by the end of the time course.

**Figure 2:**
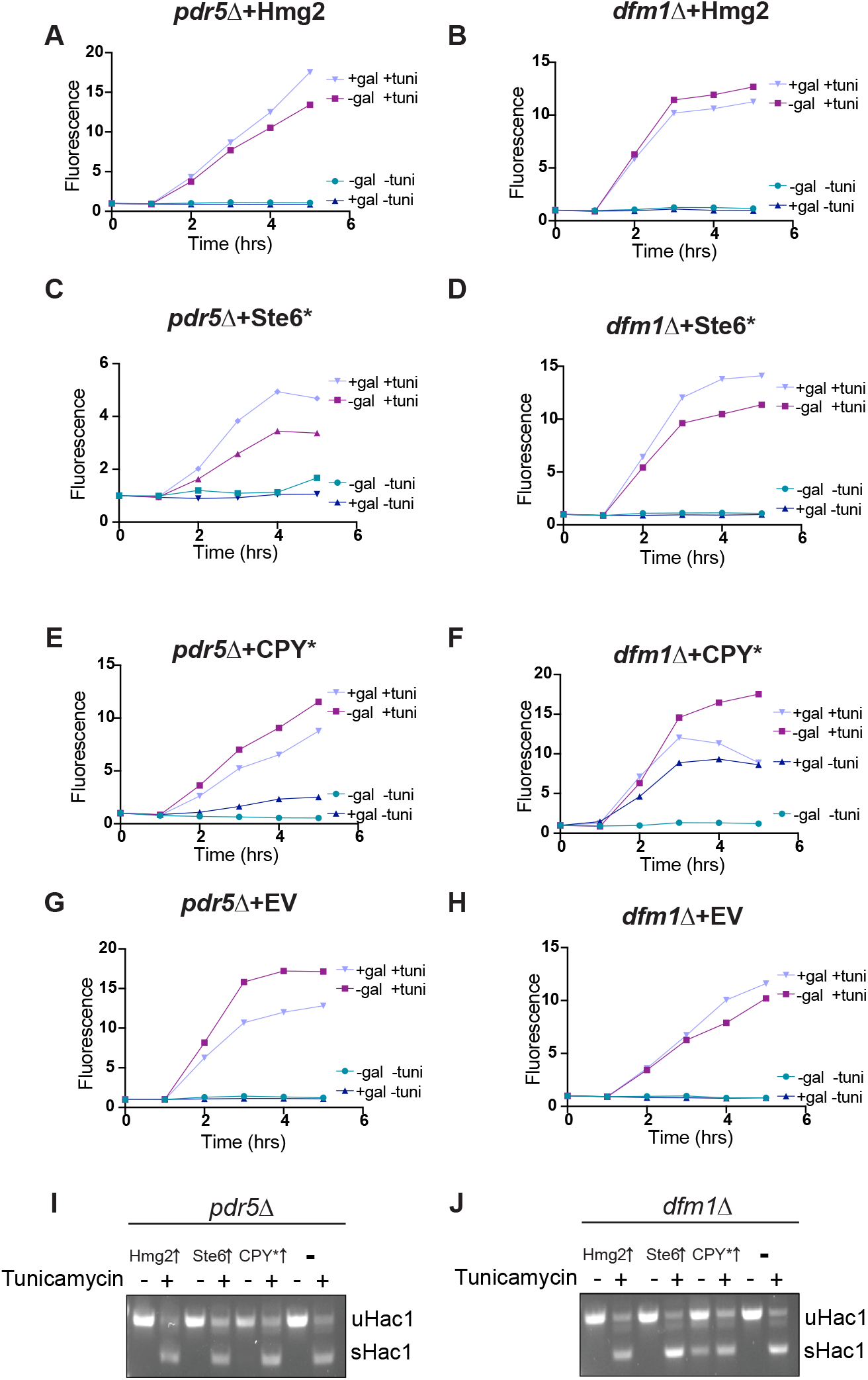
**Misfolded Membrane Protein Stress in *dfm1Δ* Cells does not Activate the Unfolded Protein Response** (A) UPR activation overtime with overexpression of a misfolded integral membrane protein. *pdr5*Δ cells containing GAL_pr_-Hmg2-6MYC and 4xUPRE-GFP (a reporter that expresses GFP with activation of the UPR) were measured for GFP expression using flow cytometry every hour for 5 hours starting at the point of galactose induction and tunicamycin or equivalent volume of DMSO was added at the 1-hour timepoint. In figure legend, +gal indicates addition of 0.2% galactose to cultures and +tuni indicates addition of 2ug/mL tunicamycin. Fluorescence is plotted as normalized fluorescence (arbitrary units) at timepoint 0-hours for each sample. (B) Flow cytometry based UPR activation assay as described in (A) except using *dfm1*Δ cells. (C) (E) and (G) Flow cytometry based UPR activation assay as described in (A) except using cells containing GAL_pr_-Ste6*-GFP, GAL_pr_-CPY*-HA, or EV, respectively. (D) (F) and (H) Flow cytometry based UPR activation assay as described in (B) except using cells containing GAL_pr_-Ste6*-GFP, GAL_pr_-CPY*-HA, or EV, respectively. (I) PCR products of spliced and unspliced Hac1 transcripts. *pdr5*Δ cells containing GAL_pr_-Hmg2-6MYC, GAL_pr_-Ste6*-GFP, GAL_pr_-CPY*-HA, or EV were treated with 0.2% galactose and 2ug/mL tunicamycin (+) or an equivalent volume of DMSO. RNA was extracted from cells and cDNA was generated and used as a template for PCR. uHac1 represents unspliced Hac1 transcripts and sHac1 represents spliced Hac1. (J) Hac1 splicing assay as in (I) except using *dfm1*Δ cells.

While CPY* exacerbates UPR activation in cells with defective ERAD-M, the inverse is not true regarding Hmg2 expression in cells with defective ERAD-L. Expression of Hmg2 did not trigger any increase above baseline UPR activation, when compared to cells containing EV, in *der1Δ* cells, where ERAD of luminal proteins is ablated (Fig. S1A&B). This indicates that inability of cells to remove misfolded membrane proteins, but not misfolded ER luminal proteins, increases sensitivity to other cellular stressors.

Our findings from flow cytometry experiments were further corroborated by measuring Hac1 splicing via polymerase chain reaction (PCR) (Fig. 2I&J). In yeast, there is one transducer of the UPR, IRE1^20^. IRE1 is a kinase and sequence-specific RNAase that when activated cleaves the mRNA of the transcription factor HAC1, creating a splice variant that is 252bp shorter and much more efficiently transcribed, resulting in more HAC1 present in the cell^21^. Activation of HAC1 by IRE1 results in the transcriptional reprogramming associated with the UPR. In our assay, samples with a band for both the spliced and unspliced variant indicated UPR activation, while a single band of the unspliced variant indicated no UPR activation. The results from these experiments were in agreement with the flow cytometry-based assay; we found no HAC1 splicing with misfolded membrane protein overexpression in *dfm1*Δ cells and an increase in HAC1 splicing in dfm1Δ cells expressing CPY* (Fig. 2J).

### Accumulation of Misfolded Membrane Proteins Upregulate Proteasome Components

After determining the UPR is not activated in *dfm1Δ* cells expressing Hmg2, we next sought to determine the transcriptional changes that occur with misfolded membrane protein stress. To address this question, we utilized RNA sequencing (RNA-seq). We prepared and sequenced cDNA libraries from mRNA extracted from *pdr5Δ* cells*, hrd1Δ pdr5Δ* cells, and *dfm1Δ pdr5Δ* cells expressing Hmg2 or EV. We used principal component analysis (PCA) to determine genes that were upregulated and downregulated most in *dfm1Δ* cells expressing Hmg2 versus the control strains; WT+EV, WT+Hmg2, *hrd1Δ*+EV, *hrd1Δ*+Hmg2, and *dfm1Δ*+EV. Principal component 1 (PC1) value of all replicate strains except for *dfm1Δ* + Hmg2 cells clustered closer to each other than they did to either replicate of the *dfm1Δ* + Hmg2 cells, indicating that these strains were transcriptionally distinct from the others sequenced (Fig. 3A).

**Figure 3:**
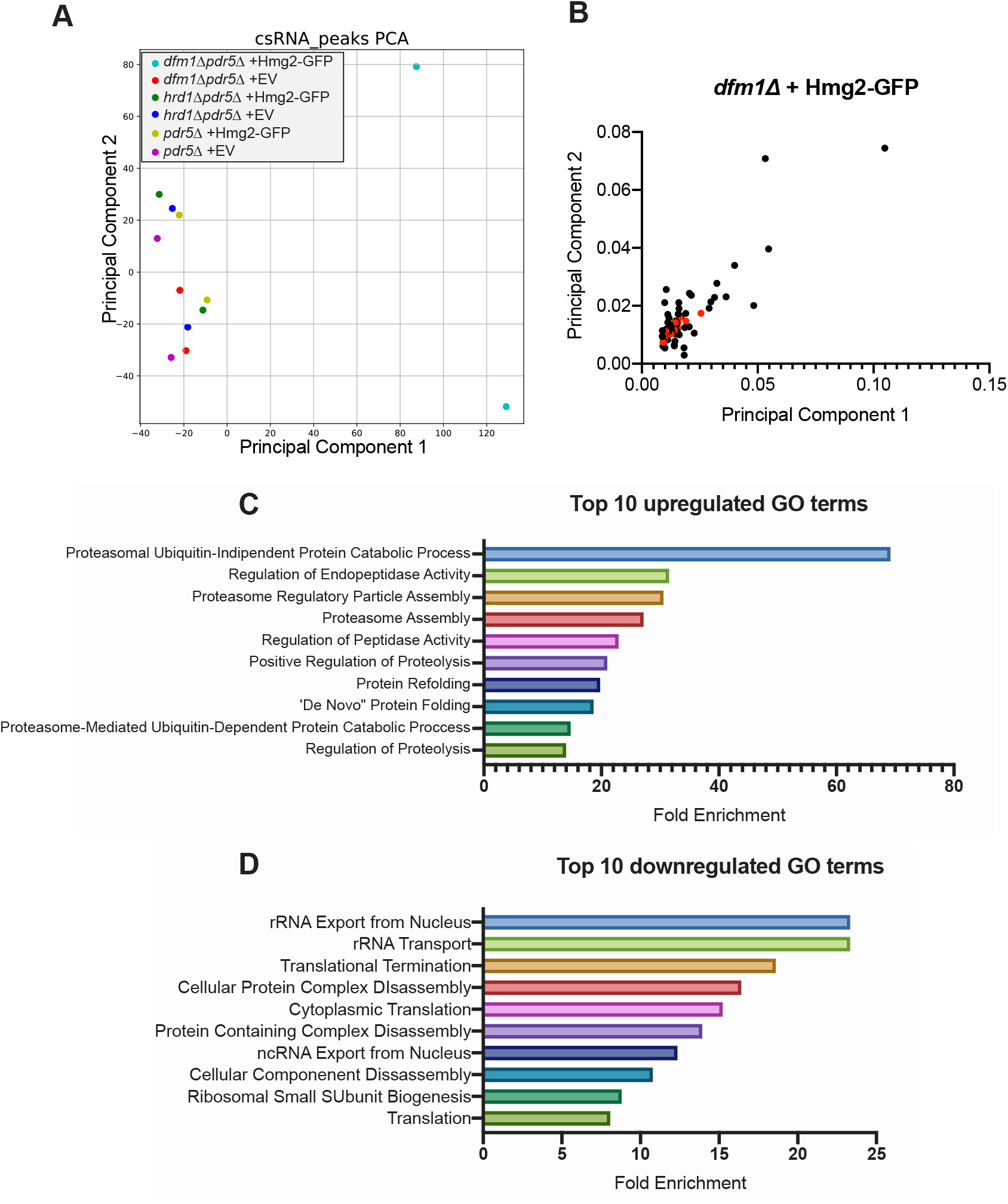
**Transcriptional Changes in Membrane Protein Stressed *dfm1Δ* Cells** (A) Principal component 1 (PC1) and principal component 2 (PC2) values of each of the two replicates of RNA-seq samples for *dfm1Δpdr5Δ*, and *hrd1Δpdr5Δ* cells containing either GAL_pr_-Hmg2-GFP or EV. (B) PC1 and PC2 of sorted top 100 highest PC1 value genes from both replicates of *dfm1Δpdr5Δ* containing GAL_pr_-Hmg2-GFP. Red dots indicate Rpn4 target genes. (C) Top 10 gene ontology (GO) terms and their enrichment factor for the set of 100 downregulated genes with the lowest PC1 scores. (D) Top 10 gene ontology (GO) terms and their enrichment factor for the set of 100 upregulated genes with the highest PC1 scores.

Upregulated (+ PC1 values) and downregulated (-PC1 values) genes in *dfm1Δ*+Hmg2 cells were used for gene ontology (GO) analysis. The most overrepresented group of upregulated genes were those classified as being involved in “Proteasomal Ubiquitin-Independent Protein Catabolic Processes”, “Regulation of Endopeptidase Activity”, and “Proteasome Regulatory Particle Assembly” (Fig. 3C). Several proteasome subunits were represented in this list of upregulated genes. The most overrepresented group of downregulated genes in this dataset were those classified as being involved in “rRNA Export from Nucleus”, “rRNA Transport”, and “Translational Termination” (Fig. 3D). Because a downregulation of the mRNA for genes encoding ribosomal proteins is a general feature of stressed yeast cells^22^, we focused on the upregulation of proteasome components. Plotting the PC1 and PC2 values for *dfm1Δ*+Hmg2 cells for the highest PC1 value genes, we observed a large overlap between genes in this dataset and those that are targets of the transcription factor Rpn4 (Fig. 3B, *highlighted in red*).

### The Transcription Factor Rpn4 is Involved in Misfolded Membrane Protein Stress

Rpn4 is a transcription factor that upregulates genes with a proteasome-associated control element (PACE) in their promoters^23^. From our RNA-seq data, there was a remarkably high overlap between the genes that were observed to be upregulated in *dfm1Δ* cells expressing Hmg2 and those that are known Rpn4 targets^23^. We reasoned that Rpn4 may be involved in adapting cells to misfolded membrane protein stress and predicted *rpn4Δ* cells should phenocopy *dfm1Δ* cells by exhibiting a growth defect induced by ERAD membrane substrates. Using the substrate-toxicity assay, we found expression of misfolded membrane proteins in *rpn4Δ* cells resulted in a growth defect equivalent to that seen in *dfm1Δ* cells (Fig. 4A&B), indicating that Rpn4 is also required for alleviating misfolded membrane protein stress. As with *dfm1Δ* cells, this effect was specific to membrane protein expression, as expression of CPY* in *rpn4Δ* cells did not result in a growth defect (Fig. 4C). This is in line with previous research demonstrating Rpn4 is activated in response to misfolded membrane protein accumulation, even in WT cells^15^. Finally, we tested a transcription factor that can regulate Rpn4, Pdr1^24^, and did not observe any growth defect in *pdr1Δ* + Hmg2 cells (Fig. S2A).

**Figure 4:**
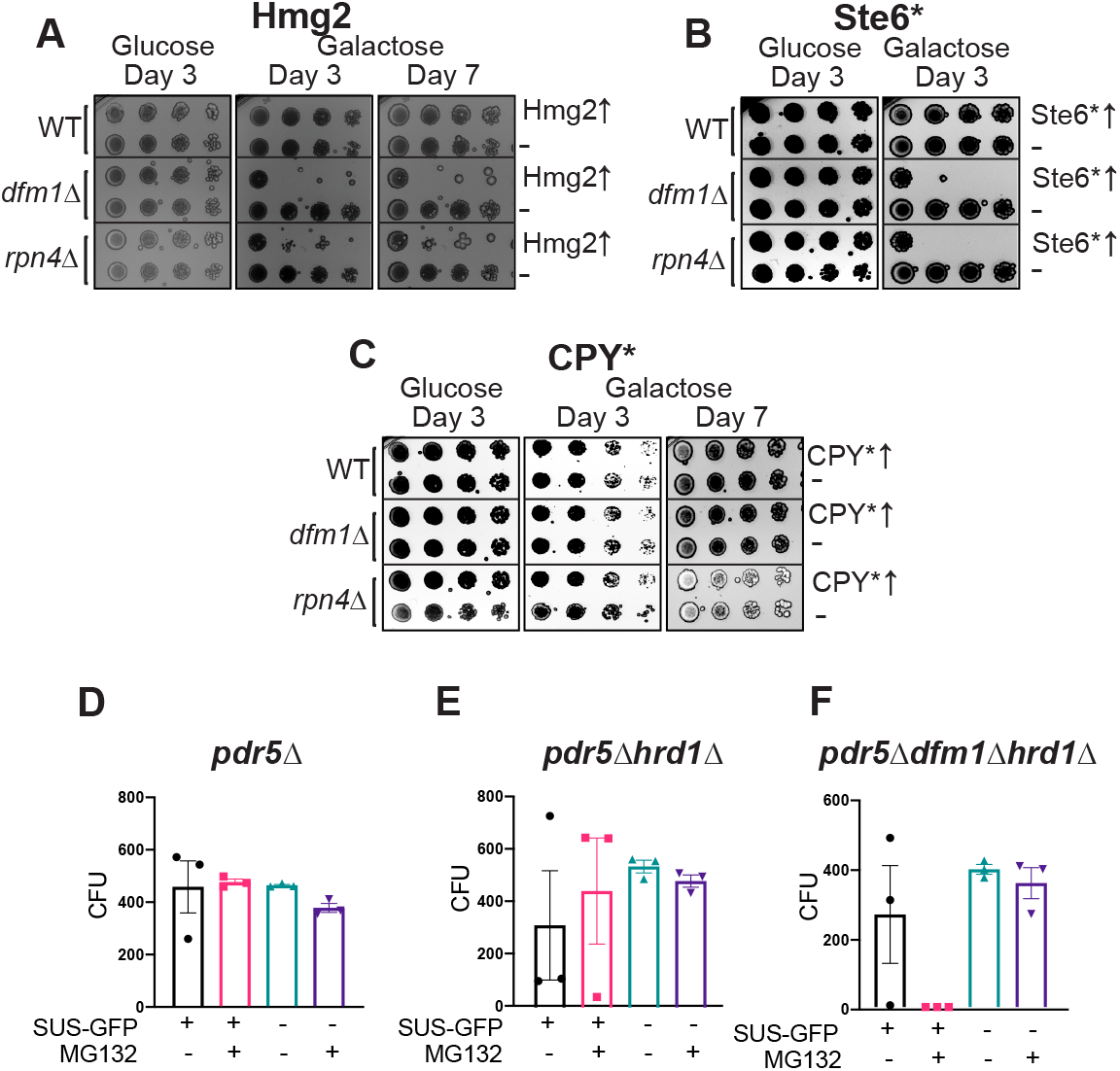
**Rpn4 is Required for Reducing Misfolded Membrane Protein Toxicity** (A) WT, *dfm1*Δ, and *rpn4Δ* cells containing either GAL_pr_-Hmg2-GFP or EV were compared for growth by dilution assay. Each strain was spotted 5-fold dilutions on glucose or galactose-containing plates to drive Hmg2-GFP overexpression, and plates were incubated at 30°C. (B) Dilution assay as depicted in (A) except using cells containing GAL_pr_-Ste6*-GFP. (C) Dilution assay as depicted in (A) except using cells containing GAL_pr_-CPY*-HA. (D) Quantification of colony forming units (CFUs) formed on appropriate selection plates from proteasome sensitivity inhibition assay. *dfm1*Δ*hrd1*Δ*pdr5*Δ cells containing SUS-GFP or EV in log phase were treated with 25uM of proteasome inhibitor MG132 or equivalent volume of DMSO for 8 hours and samples were diluted 1:500 and 50uL of each sample was plated. (E) Proteasome sensitivity assay as in (D) except using *hrd1*Δ*pdr5*Δ cells. (F) Proteasome sensitivity assay as in (D) except using *pdr5*Δ cells. For (D), (E), and (F), errors bars represent SEM. 3 biological replicates and 2 technical replicates were performed for each strain.

### Misfolded Membrane Protein Stress in dfm1Δ Cells Leads to Proteasome Impairment

Because Rpn4 appears to be active in membrane protein-stressed *dfm1Δ* cells, we hypothesized that proteasome function is impacted in *dfm1Δ* cells expressing an integral membrane protein. We tested this using an MG132 sensitivity assay developed by the Michaelis lab^15^. MG132 is a drug that reversibly inhibits proteasome function^25^. For this assay, cells in liquid culture were treated with MG132, plated, and examined for the number of colony forming units (CFUs). Due to the risk of the retrotranslocation defect being suppressed in *dfm1Δ* cells with constitutive expression of a misfolded membrane protein, and thus possibly artificially increasing the number of CFUs resulting from treatment of *dfm1Δ* cells with MG132, we opted to instead test *dfm1Δ hrd1Δ pdr5Δ* cells. These cells are unable to suppress the retrotranslocation defect of *dfm1Δ* cells, due to the absence of Hrd1, which has been characterized to function as an alternative retrotranslocon when Dfm1 is absent^13^. We utilized the engineered misfolded protein SUS-GFP. SUS-GFP contains the RING domain of Hrd1 and catalyzes its own ubiquitination, thus still causing the stress that is elicited by ubiquitinated misfolded membrane proteins in *dfm1Δ* cells^26^. We predicted that cells with compromised proteasome function will be sensitive to MG132 treatments, resulting in fewer CFUs. Strikingly, we found no CFUs upon MG132 treatment of *dfm1Δ hrd1Δ pdr5Δ* cells constitutively expressing SUS-GFP (Fig. 4F). All others strains and treatments tested did not show as dramatic of a change in the number of CFUs, either with MG132 or DMSO treatment (Fig. 4D-F). These results demonstrate that proteasome function is impacted in *dfm1Δ* cells with misfolded membrane protein accumulation.

### Ubiquitin Homeostasis is Disrupted with Misfolded Membrane Protein Accumulation

There is increasing evidence that suggests ubiquitin homeostasis and maintenance of the free ubiquitin pool is critical for cellular survival under normal and stress conditions^27–30^. Because we observed that growth defect in *dfm1Δ* cells is dependent on ubiquitination of membrane substrates, we hypothesized that ubiquitin conjugation to accumulating membrane proteins reduces the availability of free ubiquitin, impacting cell viability. Deubiquitinase (DUB) Ubp6 is a peripheral subunit of the proteasome and recycles ubiquitin from substrates prior to proteasome degradation^30^. Accordingly, *ubp6Δ* cells were employed in the substrate-toxicity assay to determine whether it is involved in alleviating misfolded membrane protein stress by replenishing the free ubiquitin pool. By utilizing the substrate-toxicity assay, we found Hmg2 or Ste6* expression causes a growth defect in *ubp6Δ* cells (Fig. 5A&B). Like *dfm1Δ* and *rpn4Δ* cells, this growth defect was specific to misfolded membrane proteins and was not observed with CPY* (Fig. 5C). To confirm whether this effect was specific to Ubp6, we also tested DUB Doa4, another regulator of free ubiquitin, in the substrate-toxicity assay. Unexpectedly, we found that *doa4Δ* cells phenocopy *ubp6Δ* cells with Hmg2 expression (Fig. 5D, Fig. S2A). From this observation, we tested a collection of DUB KOs in the substrate-toxicity assay. Of the fourteen yeast DUBs tested (out of twenty-two DUBs total), we observed a growth defect with both *ubp9Δ* and *ubp14Δ* cells (Fig. 6D, Fig. S2B). Interestingly, Ubp6, Doa4, and Ubp14 have all previously been implicated in ubiquitin homeostasis and, to date, no research has been conducted into the specific role of Ubp9^31^.

**Figure 5:**
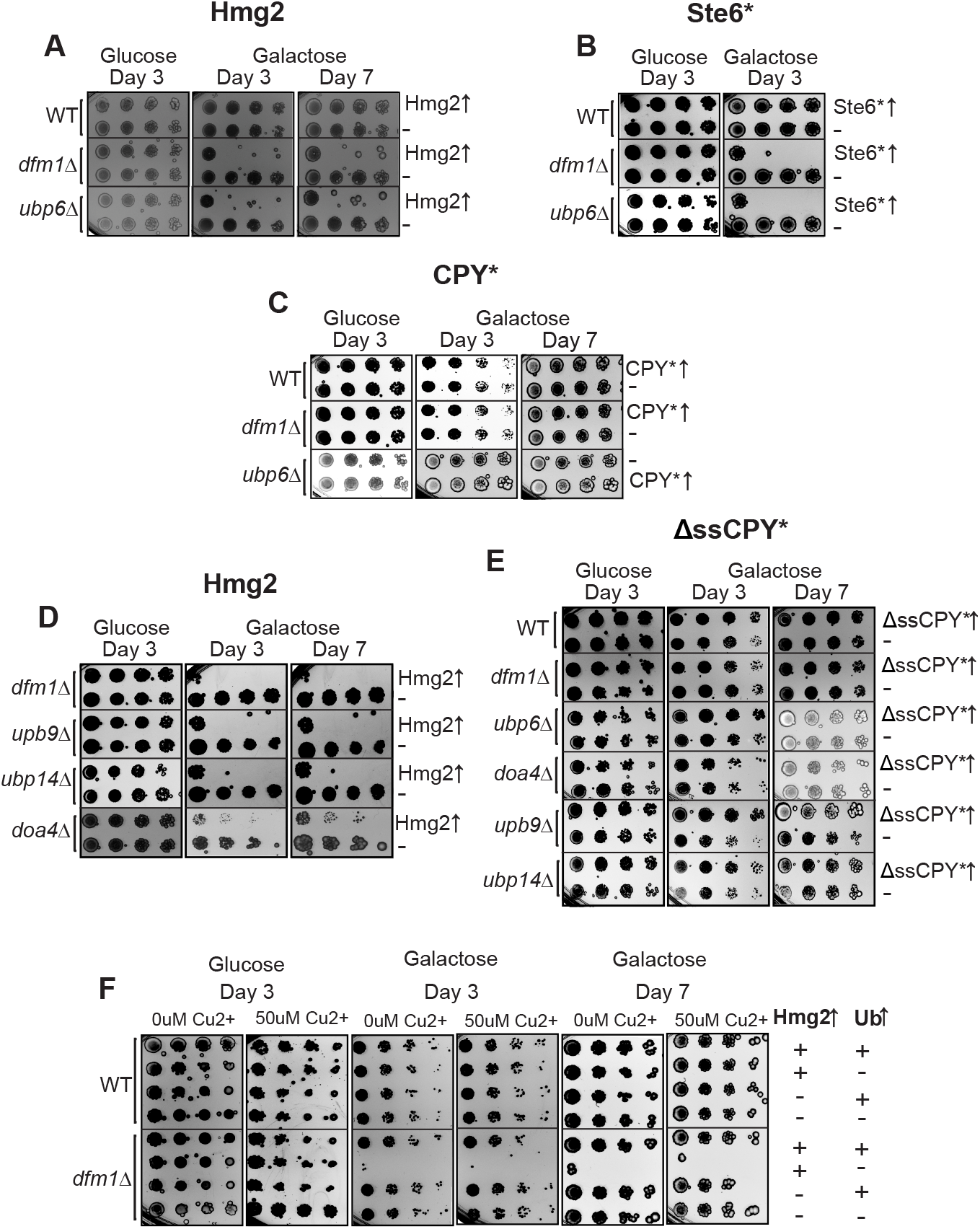
**Ubiquitin Stress Contributes to Misfolded Membrane Protein Toxicity** (A) WT, *dfm1*Δ, and *ubp6Δ* cells containing either GAL_pr_-HMG2-GFP or EV were compared for growth by dilution assay. Each strain was spotted 5-fold dilutions on glucose or galactose-containing plates to drive Hmg2-GFP overexpression, and plates were incubated at 30°C. (B) Dilution assay as in (A) except in cells containing GAL_pr_-STE6*-GFP (C) Dilution assay as in (A) except in cells containing GAL_pr_-CPY* (D) Dilution assay as described in (A) *dfm1*Δ, *ubp9Δ, ubp14Δ,* and *doa4Δ* cells. (E) WT, *dfm1*Δ, *ubp6Δ, doa4Δ, ubp9Δ,* and *ubp14Δ* cells containing either GAL_pr_-ΔssCPY*-MYC or EV were compared for growth by dilution assay. Each strain was spotted 5-fold dilutions on glucose or galactose-containing plates to drive Hmg2-GFP overexpression, and plates were incubated at 30°C. (F) WT and *dfm1*Δ cells containing either CUP1_pr_-Ub or EV were compared for growth by dilution assay. Each strain was spotted 5-fold dilutions on glucose or galactose-containing plates to drive Hmg2-GFP overexpression, and plates were incubated at 30°C. Galactose plates containing 50uM Cu2+ were used to allow expression of Ub driven by the CUP1 promoter.

**Figure 6:**
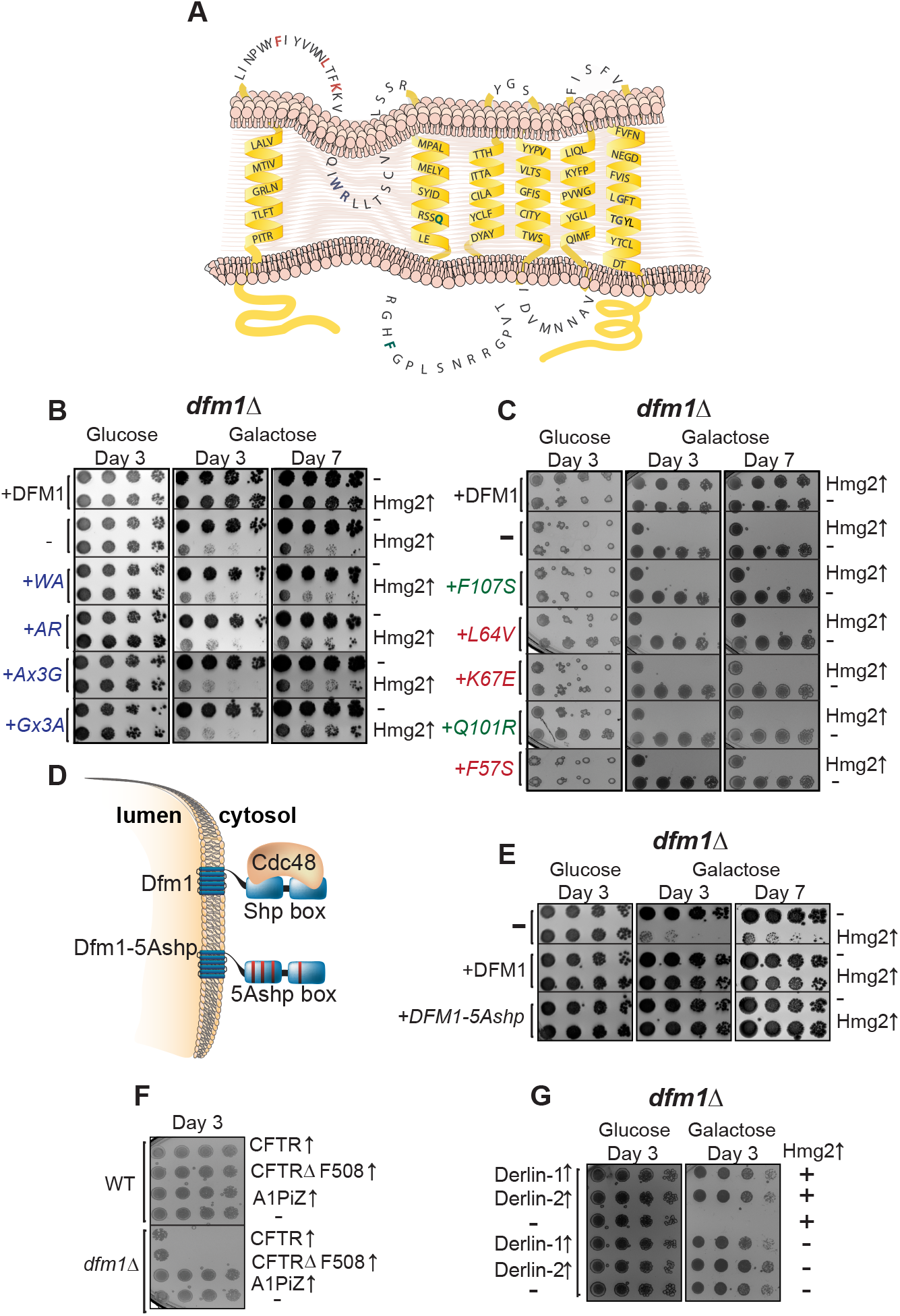
**Dfm1 Retrotranslocation Defective Mutants Show Differing Abilities to Restore Growth** (A) Depiction of Dfm1 mutants (indicated in red for L1 and green for TM 2) that have been previously identified as being retrotranslocation defective and did not restore growth in *dfm1*Δ cells expressing an integral membrane protein (GAL_pr_-Hmg2-GFP). (B) *dfm1*Δ cells with an add-back of either WT Dfm1, EV, Dfm1-WA, Dfm1-AR, Dfm1-Ax3G or Gx3A containing either GAL_pr_-Hmg2-GFP or EV were compared for growth by dilution assay. Each strain was spotted 5-fold dilutions on glucose or galactose-containing plates to drive Hmg2-GFP overexpression, and plates were incubated at 30°C. (C) Dilution assay as described in (B) except using an add-back of either WT Dfm1, EV, Dfm1(F107S), Dfm1(L64V), Dfm1(K67E), Dfm1(Q101R), or Dfm1(F57S). (D) Depiction of Dfm1 and Dfm1-5Ashp. Dfm1 is an ER-localized membrane proteins with six transmembrane domains. Both versions of Dfm1 have a cytoplasmic shp box, but the 5Ashp mutant is unable to recruit the cytosolic ATPase Cdc48. (E) Dilution assay as described in (B) except using add-back of either EV, WT Dfm1, or Dfm1-5Ashp mutant. (F) Dilution assay as described in (B) except using WT or *dfm1*Δ cells expressing human CFTR, CFTRΔ5F508, or A1PiZ. (G) Dilution Assay as described in (B) except with add-back of human Derlin-1 or Derlin-2.

One hypothesis that would explain both substrate ubiquitination dependency of the growth defect in *dfm1Δ* cells and the importance of DUBs in preventing misfolded membrane protein toxicity is that the pool of monomeric ubiquitin is depleted by accumulation of misfolded membrane proteins. If this hypothesis is correct, exogenous ubiquitin should rescue the growth defect seen from substrate-induced stress in *dfm1Δ* cells. To that end, *dfm1Δ* + Hmg2 cells harboring a plasmid containing ubiquitin under the control of the copper inducible promoter, CUP1^32^, were tested in the substrate-toxicity assay. These cells were plated on 2% galactose and 50uM copper to induce expression of Hmg2 and ubiquitin in *dfm1Δ* cells, respectively. Notably, supplementation of ubiquitin restored the growth defect (Fig. 5F). We also observed restored growth of cells plated on galactose plates without copper, which is most likely due to CUP1 being a leaky promoter^33^. This result demonstrates that increasing free ubiquitin levels can restore normal growth in these cells, indicating that levels of free ubiquitin are impacted with misfolded membrane protein stress.

If ubiquitin depletion was the only disruption to ubiquitin homeostasis in all the DUB KOs tested, it would be expected that expression of any ubiquitinated misfolded protein would cause growth stress. We expressed the cytosolic misfolded protein ΔssCPY*, which is not targeted by cytosolic protein quality control machinery (not by ERAD)^34^, in *ubp6Δ, doa4Δ, ubp9Δ,* and *ubp14Δ* cells in the substrate-toxicity assay (Fig. 5E). ΔssCPY* expression did not elicit growth stress in any of the strains tested (Fig. 5E). This finding implies misfolded membrane proteins alter ubiquitin homeostasis differently than cytosolic proteins, or that Ubp6, Doa4, Ubp14, and Ubp9 prevent misfolded membrane protein toxicity through a different mechanism than directly affecting the abundance of monomeric ubiquitin.

### Deubiquitinases and RPN4 Function in the Same Pathway as DFM1, but Not in Parallel with Each Other

We tested double knockouts of *dfm1Δ rpn4Δ, dfm1Δ ubp6Δ,* and *rpn4Δ ubp6Δ* cells in the substrate-toxicity assay to determine whether these genetic components function within the same or parallel pathways. Expression of either Hmg2 or Ste6* in either *dfm1Δrpn4Δ* or *dfm1Δubp6Δ* cells resulted in a growth defect that phenocopied that observed in any of the single knockouts (Fig. S3A&B), whereas expression of CPY* showed no growth defect (Fig. S3C). In contrast, double-null *rpn4Δubp6Δ* cells showed a growth defect in the absence of substrates whereas *rpn4Δ* and *ubp6Δ* displayed normal growth. Moreover, *rpn4Δubp6Δ* cells along with expression of Hmg2 or Ste6* resulted in synthetic lethality (Fig. S3A-C), further indicating Rpn4 and Ubp6 act in parallel in alleviating membrane protein toxicity.

We also tested expression of previously described Hmg2 mutants K6R and K357R in *rpn4Δ* and *ubp6Δ* cells (Fig. S3E). As with *dfm1Δ* cells expressing these mutants, expression of Hmg2-K6R does not cause toxicity while Hmg2-K357R does cause toxicity in both *rpn4Δ* and *ubp6Δ.* Thus, ubiquitination of misfolded membrane proteins influences toxicity in *dfm1Δ, rpn4Δ,* and *ubp6Δ* cells.

### Increased Expression of Dfm1 Relieves Misfolded Membrane Protein Stress in rpn4Δ and ubp6Δ Cells

Using the substrate-toxicity assay, we examined whether increasing expression of Dfm1 could relieve growth stress in *rpn4Δ* and *ubp6Δ* cells expressing Hmg2. We utilized the substrate-toxicity assay with the addition of galactose inducible Dfm1 to address this question. Increasing Dfm1 in both *rpn4Δ+*Hmg2 and *ubp6Δ*+Hmg2 cells restored normal growth (Fig. S3D). Importantly, endogenous Dfm1 is already present in these cells, but increasing expression level relieves toxicity caused by misfolded membrane proteins.

### Dfm1 has a Dual Role in ER Protein Stress and ERAD Retrotranslocation

Previous work from the Hampton lab establishing a role for Dfm1 in misfolded membrane protein retrotranslocation also identified several motifs of Dfm1 that are essential for its retrotranslocation function^7^. Additionally, by employing an unbiased genetic screen, recent work from our lab identified five residues of Dfm1 that are required for retrotranslocation^35^. Here, we tested whether these residues, critical for Dfm1’s retrotranslocation function, are required for alleviating the growth stress in *dfm1Δ* cells expressing Hmg2.

Fig. 6A shows the amino acid sequence of Dfm1 as well as its predicted topology, spanning seven transmembrane, with the highlighted motifs and residues that were previously characterized to be critical for Dfm1’s ERAD function^7, 35^. Dfm1 contains two motifs that are well conserved amongst the rhomboid superfamily, the WR motif in Loop 1 and the GxxxG (Gx3G) motif in transmembrane domain (TMD) 6^36^. Both of these motifs are required for Dfm1-mediated retrotranslocation^7, 37^. We first tested the requirement of the conserved rhomboid motif mutants by expressing Hmg2 with WR mutants (WA and AR) and Gx3G mutants (Ax3G and Gx3A) and observed no restoration in growth (Fig. 6B). Our previous work determined that Loop 1 mutants (F58S, L64V, and K67E) obliterated Dfm1’s ability to bind a subset of misfolded membrane substrates, and TMD 2 mutants (Q101R and F107S) reduce the lipid thinning ability of Dfm1, a function which aids in Dfm1’s retrotranslocation function^37^. Accordingly, we utilized these mutants in our growth assay and did not observe a rescue of the growth defect (Fig. 6C). We have previously shown that alteration of the five signature residues of the Dfm1 SHP box to alanine (Dfm1-5Ashp) ablates its ability to recruit Cdc48 (Fig. 6D). We also established, Dfm1’s Cdc48 recruitment function is required for Dfm1’s retrotranslocation function, whereas the Dfm1-5Ashp mutant impairs its retrotranslocation function^7^. Notably, in contrast to the other mutants tested, Dfm1-5Ashp was still able to alleviate the growth defect the same as WT Dfm1 (Fig. 6E). These results suggest that Dfm1’s substrate engagement and lipid thinning function is required for alleviating membrane substrate-induced stress whereas Dfm1’s Cdc48 recruitment function is dispensable for alleviating the growth stress.

### Human Membrane Protein Causes Growth Stress and Human Derlins Relieve Growth

*Stress* Since a wide variety of misfolded membrane proteins elicit growth stress in *dfm1Δ* cells, we hypothesized that growth stress would also be observed with expression of clinically relevant human misfolded membrane proteins. We tested expression of WT cystic fibrosis transmembrane receptor (CFTR), CFTRΔF508, the most common disease-causing variant of CFTR, and the Z variant of alpha-1 proteinase inhibitor (A1PiZ), a protein variant that results in alpha-1 antitrypsin deficiency (AATD). CFTR and CFTRΔF508 are ERAD-M substrates when expressed in yeast, while A1PiZ is a soluble misfolded protein targeted by ERAD-L^38, 39^. When these proteins were expressed in *dfm1Δ* cells, both CFTR and CFTRΔF508 resulted in a growth defect, while none was observed with expression on A1PiZ (Fig. 6F). Expression of any of the proteins in WT yeast cells resulted in no growth defect. While we had originally hypothesized that only CFTRΔF508 would cause a growth defect when expressed in *dfm1Δ* cells, it was not wholly surprising that WT CFTR also elicited growth stress. Previous studies have shown that while virtually all CFTRΔF508 is targeted to ERAD, about 80% of WT CFTR is degraded via ERAD in yeast and mammals^38, 40, 41^.

Dfm1 is a rhomboid pseudoprotease, and a member of the derlin subclass of rhomboid proteins^37^. The human genome encodes three derlins, Derlin-1, Derlin-2, and Derlin-3. Yeast Dfm1 is the closest homolog of the mammalian derlins^36^. All three are ER localized proteins that are implicated in ERAD and adaptation to ER stress^42–47^. We expressed human Derlin-1 and Derlin-2 in *dfm1Δ*+Hmg2 cells. Both human derlins were able to rescue growth in these cells in the substrate-toxicity assay (Fig. 6G). This was surprising, as we had previously found that mammalian derlins cannot complement the retrotranslocation function of Dfm1 in yeast cells for self-ubiquitinating substrate (SUS)-GFP, a similar substrate to Hmg2^35^.

### Dfm1 Solubilizes Misfolded Membrane Protein Aggregates Independent of Cdc48 Recruitment

The above studies show that Dfm1 residues critical for retrotranslocation —through substrate binding and its lipid thinning function— are also important for alleviating membrane substrate-induced stress. Conversely, Dfm1’s Shp box mutant (Dfm1-5Ashp) (Fig. 6D) that does not recruit Cdc48 exhibits normal growth in the substrate-toxicity assay. We surmise that Dfm1’s actions —independent of its Cdc48 recruitment function— may be directly acting on misfolded membrane substrates to prevent growth stress. One possibility is that Dfm1 may directly act on misfolded membrane substrates by functioning as a chaperone. We hypothesize that Dfm1 acts as either a holdase, preventing the aggregation of misfolded membrane substrates, or as a disaggregase, disaggregating existing protein aggregates. This function of Dfm1 would allow it to prevent cellular stress from misfolded membrane proteins. To address this hypothesis, fluorescence microscopy was used on *dfm1Δ* cells containing constitutively expressed Hmg2-GFP to visualize membrane substrate aggregation. Indeed, fluorescent microscopy images and subsequent analysis revealed the majority of Hmg2-GFP in *dfm1Δ* cells form fluorescent puncta (Fig. 7C). Moreover, both wildtype Dfm1 and SHP mutant Dfm1-5Ashp add-back in *dfm1Δ* +Hmg2 cells showed a significant reduction in the fraction of Hmg2-GFP in puncta. To confirm the fluorescent puncta are aggregates, we employed a detergent solubility assay. Microsomes were isolated and incubated in dodecyl maltoside (DDM) and subjected to centrifugation to separate aggregated substrate (pellet fraction) from solubilized substrate (supernatant fraction). As shown in Fig. 7A, only a small amount of Hmg2-GFP in *dfm1Δ* cells was soluble. Conversely, with Dfm1 and Dfm1-5Ashp add back cells, there was a significant increase in detergent-solubilized Hmg2-GFP. As a control for these studies, we tested a properly folded ER membrane protein, Sec61-GFP. In contrast to Hmg2-GFP, majority of Sec61-GFP was in the detergent-solubilized supernatant fraction and there was no change in Sec61-GFP detergent solubility with Dfm1 or Dfm1-5Ashp addback (Fig. S4A). It appears Dfm1— independent of its Cdc48 recruitment function—functions as a holdase and prevents the aggregation of misfolded membrane proteins. We next explored additional Dfm1 residues that are required for solubilizing membrane substrates. Accordingly, mutants in the conserved rhomboid mutants (AR and Ax3G), were employed in the solubility assay and all mutants tested were unable to promote Hmg2 solubilization when expressed in *dfm1Δ* cells (Fig. 7A&B). This was supported through visualization of Hmg2-GFP puncta in the Dfm1-AR and Dfm1-Ax3G add-backs (Fig. 7C&D). Altogether, with all criteria examined, Dfm1 is critical influencing the solubility of its ERAD membrane substrate (Fig. 7E). Although Dfm1’s conserved rhomboid motifs are critical for its role preventing cellular stress, its Cdc48 recruitment activity – critical for retrotranslocation--is not required for this newly-established chaperone function.

**Figure 7:**
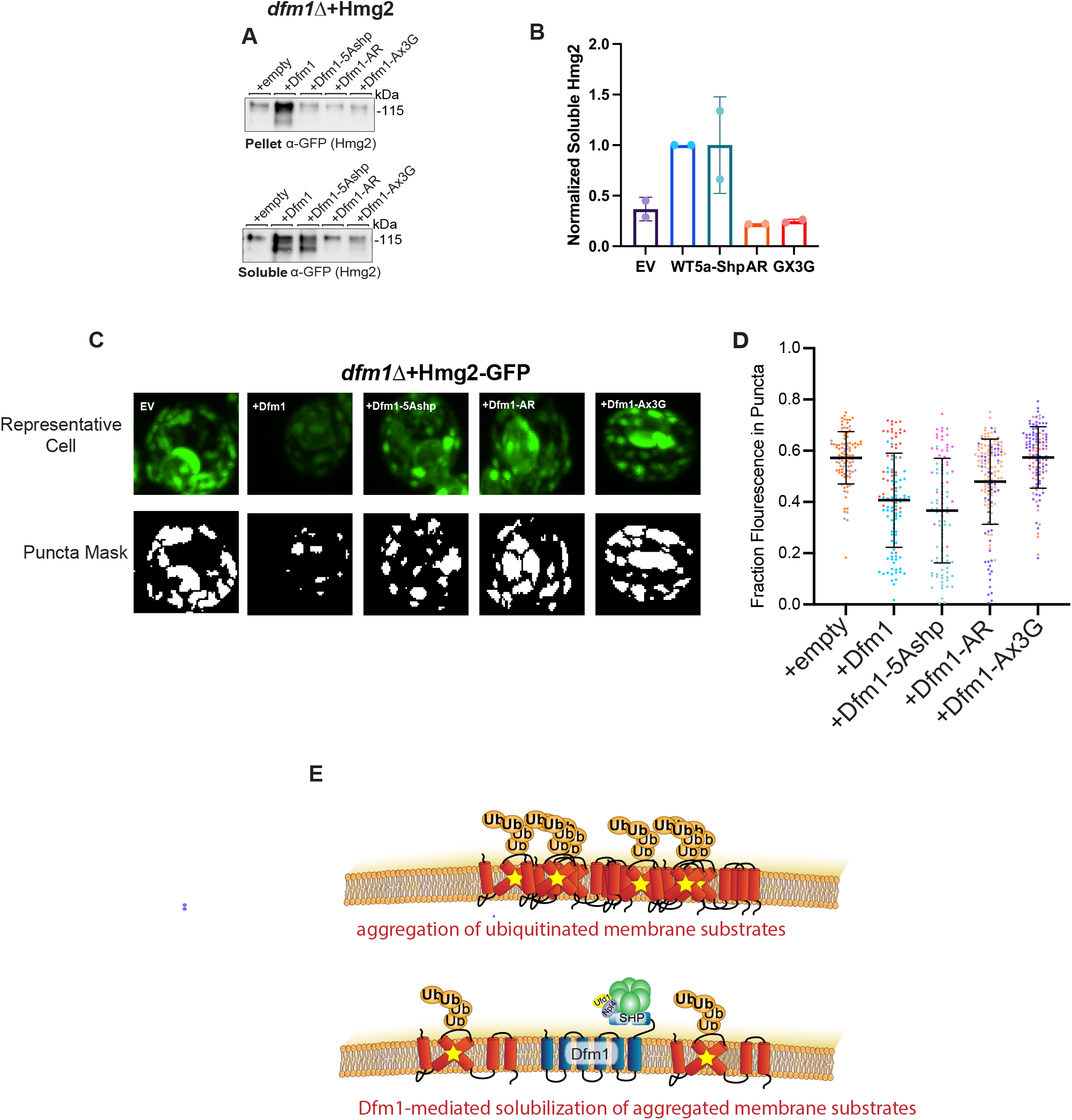
**Dfm1 Reduces Misfolded Membrane Protein Toxicity by Acting as an ATP-Independent Disaggregase** (A) Western blot of aggregated versus soluble membrane proteins at the ER. Lysates from *dfm1*Δ cells containing HMG2-GFP, with either add-back of EV, WT DFM1, DFM1-5Ashp, Dfm1-AR, Dfm1-AxxxG, Dfm1-L64V, Dfm1-F107S, Dfm1-K67E, Dfm1-Q101R, and Dfm1-F58S were blotted using anti-GFP to detect HMG2. Top: ER aggregated fraction. Bottom: ER soluble fraction. (B) Quantification of band intensity of soluble Hmg2 in (A). Intensity was normalized to soluble Hmg2 intensity in *dfm1*Δ cell containing Hmg2 with add-back of WT Dfm1. Errors bars represent standard error of the mean for 2 biological replicates. (C) Representative confocal microscopy images and puncta mask of Hmg2-GFP in *dfm1*Δ cells with add-back of EV, WT DFM1, DFM1-5Ashp, Dfm1-AR, and Dfm1-AxxxG. Two biological replicates were imaged, and two images were taken of each strain. (D) Fraction of Hmg2-GFP in puncta for *dfm1*Δ cells with add-back of EV, WT DFM1, DFM1-5Ashp, Dfm1-AR, and Dfm1-AxxxG. Each dot represents an individual cell and each color of dots for each strain indicates the specific biological replicate. Error bars represent SEM. (D) Model depicting integrated model of Dfm1’s function in misfolded membrane protein stress. Top: Misfolded membrane proteins in the absence of Dfm1 forming aggregates within the ER membrane. Bottom: Cells with WT Dfm1 or 5Ashp-Dfm1 disaggregating misfolded membrane proteins and preventing cellular toxicity.

## DISCUSSION

Proper protein folding and efficient elimination of misfolded proteins is imperative for maintaining cellular health. Accumulation of misfolded proteins, which is a widespread phenomenon in aging and diseased cells, is deleterious to cells and can impact cellular function. Despite membrane proteins accounting for one-third of proteins in the cell, there is a dearth of research on the source of stress that is triggered by accumulation of misfolded membrane proteins. In this study, we sought to understand how cells are impacted by misfolded membrane protein stress and how they prevent toxicity from these misfolded proteins. By employing transcriptomic analyses and our genetically tractable substrate-toxicity assay, we found that the source of cell toxicity was from aggregation of accumulated misfolded membrane proteins and that Dfm1’s rhomboid motifs are required for solubilizing aggregation-prone substrates. We propose a model in which *dfm1Δ* cells form aggregates of ubiquitinated ERAD membrane substrates, which compromises proteasome function and disrupts ubiquitin homeostasis. Overall, our studies unveil a new role for rhomboid pseudoproteases in mitigating the stress state caused by ERAD membrane substrates, a function that is independent of their retrotranslocation function.

Our results above (Fig. 6B-D) indicate differential requirements for Dfm1’s role in membrane substrate retrotranslocation, versus its role in stress alleviation. These results are fascinating, because of all the retrotranslocation-deficient mutants tested, we were able to identify a mutant that was still able to rescue the growth defect observed in *dfm1Δ*+Hmg2 cell. This indicates a bifurcated role of Dfm1 in retrotranslocation and membrane protein stress alleviation. The retrotranslocation defective mutants that did not restore growth were mutations of conserved rhomboid protein motifs (WR and Gx3G), mutants that obliterate substrate engagement (Loop 1 mutants: F58S, L64V, and K67E), and mutants that reduce the ability of Dfm1 to distort the ER membrane (TMD 2 mutants: Q101R and F107S). This indicates the substrate binding and lipid distortion roles of Dfm1 that are imperative for retrotranslocation are also imperative for alleviation of misfolded membrane protein stress. In contrast, the SHP box mutant, which prevents Cdc48 binding to Dfm1, restores growth in *dfm1Δ* + Hmg2 cells (Fig. 6D). While Cdc48 binding to Dfm1 is critical for retrotranslocation, this is not a requirement for Dfm1’s role in preventing membrane proteotoxicity. Previous work from our lab indicates transient interactions between membrane substrates and Dfm1 still occurs even when Dfm1’s Cdc48 recruitment activity is impaired^35^. This suggests that this level of physical interaction is sufficient for Dfm1 to directly act on substrates to prevent membrane substrate-induced stress.

We previously demonstrated that expression of integral membrane ERAD substrate induces toxicity in yeast cells when Dfm1 function is impaired^16^. Remarkably, this strong growth defect phenotype is unique to *dfm1Δ* strains: other equally strong ERAD deficient mutants, both upstream or downstream of Dfm1 (*hrd1*Δ or *cdc48-2),* show no growth stress upon similar elevation of ERAD integral membrane substrates. Thus, the growth effects above suggest the intriguing possibility that Dfm1 has a unique role in this novel ER stress. By analyzing the transcriptome upon triggering this unique membrane substrate-induced stress state, we find that many proteasomal subunits are upregulated. Interestingly, Rpn4 – a transcription factor known to induce proteasome subunit expression – upregulates many of the proteasomal subunits upregulated in our transcriptome analysis. One interpretation of our data is that accumulation of integral membrane proteins results in reduced proteasome efficiency, which triggers Rpn4-mediated upregulation of proteasome subunits. Indeed, we and others have shown that *rpn4Δ* cells phenocopy *dfm1Δ* cells by exhibiting a growth defect upon expression of ER integral substrates, and not ERAD-L substrates^15^. This was also supported by our above studies showing ERAD-M substrates exacerbate cellular growth defects when proteasome function is compromised with treatment of proteasome inhibitor, MG132 (Fig. 4D-F). These data indicate that cells require optimal proteasome activity to avoid the proteotoxicity associated with integral membrane ERAD substrates.

The facile and genetically tractable substrate-toxicity assay allowed us to ascertain how membrane substrates cause the growth defect phenotype when Dfm1 is absent. Intriguingly, no growth defect was observed in *dfm1Δ* expressing the K6R Hmg2 mutant (with negligible ubiquitination), while the K357R Hmg2 mutant (with slight ubiquitination) still showed a growth defect, suggesting the source of Dfm1-mitigated stress is ubiquitination of the substrates. We reasoned that accumulation of ubiquitinated ERAD membrane substrates disrupts the ubiquitin pool through excessive ubiquitination of substrates and concomitant depletion of the ubiquitin pool. Indeed, a collection of DUB mutants (*ubp6*Δ, *doa4*Δ, *ubp14*Δ) --known for their role in replenishing the ubiquitin pool through their deubiquitinating function -- is unable to mitigate the proteotoxic effect of integral membrane substrates and proteotoxic stress is rescued with exogenous addition of ubiquitin molecules in *dfm1Δ*+Hmg2 cells. This observation is extended in mammalian studies in which a mouse line with a loss-of-function mutation in Usp14, the mammalian homolog of Ubp6, reduction in the pool of free ubiquitin in neurons results in ataxia that can be rescued with exogenous ubiquitin expression^28^. Perhaps what is most fascinating is that the stress state is only induced by excessive ubiquitination of integral membrane substrates and not soluble proteins residing in the ER lumen or the cytosol, suggesting the source of stress is due to excessive ubiquitination of substrates at the ER membrane. One possibility is that ubiquitination of membrane proteins not only depletes the pool of cellular ubiquitin, but also increases the propensity of membrane proteins to aggregate. This hypothesis will need to be explored in future studies.

Our data on the ability of Dfm1 to influence misfolded membrane protein solubility provides evidence that this is the mechanism by which Dfm1 prevents misfolded membrane protein toxicity. We find that both WT Dfm1 and Dfm1-5Ashp promote solubility of Hmg2 (Fig. 7A-C). In contrast, the retrotranslocation defective Dfm1 rhomboid motif mutants, Dfm1-AR and Dfm1-Ax3G are not able to promote Hmg2-GFP solubility. This is in agreement with our observation that both WT Dfm1 and Dfm1-5Ashp can restore normal growth in *dfm1Δ* cells in the S-T assay, but the rhomboid motifs mutants cannot (Fig. 6B&D). The exact mechanism by which Dfm1 influences Hmg2 solubility is unclear. We propose two possible models that will be important to distinguish between in future works. In one model, Dfm1 functions as a disaggregase to physically separate misfolded membrane proteins from existing protein aggregates. In another model, Dfm1 functions as a holdase to promote solubility of misfolded membrane proteins and limit their ability to aggregate. While the ability of Dfm1-5Ashp to increase Hmg2 solubility in *dfm1Δ* cells indicates that Dfm1’s chaperone ability is ATP-independent, we cannot exclude the possibility that Dfm1 recruits another ATPase besides Cdc48, independent of the SHP box motif.

Another possibility is that Dfm1 itself can bind and hydrolyze ATP. There are a growing number of identified ATP-independent disaggregases^48^, including one membrane protein dissagregase identified in plants^49^. Understanding how Dfm1 influences the solubility of membrane substrates will be an important future line of inquiry.

Protein aggregation has been linked to many human maladies, including neurodegenerative disorders like Parkinson’s and Alzheimer’s disease. While protein aggregates are commonly recognized as a feature of disease, the exact mechanism by which they prove toxic to cells is elusive in many cases. Protein aggregation has been shown to induce toxicity through sequestration of cellular components^50^. In our system, it is possible that aggregated membrane proteins sequester ubiquitin, thereby disrupting ubiquitin homeostasis, as well as sequestering proteasomes. It will be interesting to determine the protein composition of ER aggregates in the absence of Dfm1. Nevertheless, this toxicity is prevented by Dfm1’s chaperone function. One possible model for Dfm1’s function is that ERAD substrates are solubilized during the retrotranslocation process, relying on Dfm1’s rhomboid motifs, substrate engagement function, and lipid thinning function.

While the structure of Dfm1 has not been determined, there is a structure of its mammalian homolog, Derlin-1^51^. The cryo-EM structure of Derlin-1 reveals that it forms a homotetramer with a pore in the center. If this is the type of structure derlins form to retrotranslocate proteins, a chaperone ability would be necessary to ensure that substrates were properly solubilized to pass through the pore. Previous work from the Brodsky lab demonstrated that aggregation-prone ER proteins are more likely to be targeted by ERAD and are disaggregated by the ATP-dependent cytoplasmic disaggregase Hsp104, which aids in retrotranslocation^52^. Our results demonstrate that a component of membrane protein retrotranslocation machinery, Dfm1, also has a chaperone function to aid in retrotranslocation.

Molecular chaperones have long been identified for their role in protein quality control systems, including ERAD, for their ability to triage terminally misfolded proteins to degradation machinery. In recent years, more studies have shown a dual function of protein quality control machinery in directly controlling degradation and being chaperones^53, 54^. We have now provided evidence for rhomboid pseudoproteases, a subclass of proteins widely recognized as involved in protein quality control, having chaperone function. This raises the question of whether chaperone ability is more widespread among other protein quality control components, specifically those known to bind to membrane proteins. The Carvalho group has demonstrated that the Asi complex involved in inner nuclear membrane protein quality control in yeast and the mammalian ERAD factor membralin are able to recognize transmembrane domains of misfolded proteins^55, 56^. It is possible that chaperone function has arisen more than once evolutionarily among proteins involved in membrane protein quality control.

Rhomboid pseudoproteases have been recognized for over a decade as being involved in a diverse array of cellular process, from protein quality control to cell signaling to adaptations to cellular stress^57–61^. Our lab and others have made progress towards understanding how these proteins are able to function is such diverse cellular process without an enzymatic function. With the knowledge that Dfm1 is an ATP-independent chaperone, it will be of extreme interest to determine if this function is conserved among all rhomboid pseudoproteases, and even among the active rhomboid proteases. Two specific areas of interest include determining the conservation of this chaperone function and identifying the repertoire of substrates that can be solubilized by rhomboid pseudoproteases. There are two subclasses of rhomboid pseudoproteases, iRhoms and derlins. Both of these classes are evolutionarily distinct and it will be of interest to determine if chaperone ability is only specific to derlins, and not to iRhoms^62^. Derlins are known to function in retrotranslocation of a wide variety of substrates, including disease-associated membrane substrates. In this study, we observed accumulation of both WT and the disease causing CFTRΔF508 caused growth stress in *dfm1Δ* cells^11, 36, 42, 61^. Surprisingly, we found that heterologous expression of both human Derlin-1 and Derlin-2 restores growth in yeast *dfm1Δ*+Hmg2 cell implying the solubility function is a conserved feature amongst all derlin rhomboid pseudoproteases. Moreover, research from our lab demonstrated that Derlin-1 and Derlin-2 does not support ERAD-M retrotranslocation in *dfm1Δ* cells^35^. This indicates that Derlin-1 and Derlin-2 relieves toxicity in *dfm1Δ* + Hmg2 cells, without restoring retrotranslocation, likely by a conserved chaperone function.

Our studies provide the first evidence that the derlin subclass of rhomboid pseudoproteases function as chaperones by influencing the solubilization of misfolded membrane substrates. Findings gleaned from our studies hold great promise for foundational and translational arenas of cell biology, since fundamental understanding of a membrane protein chaperone will aid in understanding a plethora of diseases associated with misfolded membrane proteins such as cystic fibrosis, retinal degeneration, and neurodegenerative diseases.

## AUTHOR CONTRIBUTIONS

R.K. and S.E.N. designed research; R.K., J.J., D.S., T.K., A.A., S.D., S.L., and S.E.N. performed research; R.K., D.S., J.J., T.K., S.D., L.S., and S.E.N. analyzed data; R.K. and S.E.N. wrote the paper; and R.K. and S.E.N. designed illustrations for figures. All authors reviewed the results and approved the final version of the manuscript.

## ACKNOWLEDGEMENTS

We thank Tom Rapoport (Harvard Medical School), Davis Ng (National University of Singapore), Randy Schekman (University of California, Berkeley), and Susan Michaelis (John Hopkins University), and Jeff Brodsky (University of Pittsburgh) for providing plasmids and antibodies. We also thank the Neal lab members for their positive reinforcement, in depth discussions and technical assistance. These studies were supported by NIH grant 1R35GM133565-01, Pew Biomedical Award, and NSF CAREER grant to S.E.N. S.D.H. is supported by NIH grant K99GM135515.

## RESOURCE AVAILABILITY

### Lead contact

Further information and requests for resources and reagents should be directed to and will be fulfilled by the Lead Contact, Sonya Neal.

### Materials availability

Plasmids and yeast strains generated in this study is available from our laboratory.

### Data and Code Availability

Code for microscopy puncta analysis will be made available.

---

**Figure S1: UPR Activation in *der1*Δ cells**

A. UPR activation overtime with overexpression of a misfolded integral membrane protein. der1Δ cells containing GALpr-Hmg2-6MYC and 4xUPRE-GFP (a reporter that expresses GFP with activation of the UPR) were measured for GFP expression using flow cytometry every hour for 5 hours starting at the point of galactose induction and tunicamycin or equivalent volume of DMSO was added at the 1-hour timepoint. In figure legend, +gal indicates addition of 0.2% galactose to cultures and +tuni indicates addition of 2ug/mL tunicamycin. Fluorescence is plotted as normalized fluorescence (arbitrary units) at timepoint 0-hours for each sample.
B. Flow cytometry based UPR activation assay as described in (A) except with cells containing EV.

**Figure S2: Misfolded Membrane Proteins do not Elicit Stress in cells lacking Transcription Factor Pdr1 or Most Deubiquitinases**

A. WT, dfm1Δ, doa4Δ, and pdr1Δ cells containing either GALpr-HMG2-GFP or EV were compared for growth by dilution assay. Each strain was spotted 5-fold dilutions on glucose or galactose-containing plates to drive Hmg2-GFP overexpression, and plates were incubated at 30°C.
B. Dilution assay as in (A) except in dfm1Δ, ubp2Δ, ubp5Δ, miy1Δ, ubp8Δ, miy2Δ, otu2Δ, ubp1Δ, ubp11Δ, ubp7Δ, and ubp3Δ cells.

**Figure S3: Genetic Interactions Between Dfm1, Rpn4, and Ubp6 in Resolving Misfolded Membrane Protein Toxicity**

A. dfm1Δ, dfm1Δrpn4Δ, dfm1Δubp6Δ, and rpn4Δubp6Δ cells containing either GALpr-HMG2-GFP or EV were compared for growth by dilution assay. Each strain was spotted 5-fold dilutions on glucose or galactose-containing plates to drive Hmg2-GFP overexpression, and plates were incubated at 30°C.
B. Dilution Assays as depicted in (A) except using cells containing GALpr-STE6*-GFP.
C. Dilution Assays as depicted in (A) except using cells containing GALpr-CPY*.
D. dfm1Δ, rpn4Δ, and ubp6Δ cells containing either GALpr-Hmg2-GFP or EV and GALpr-Dfm1-10xHis or EV were compared for growth by dilution assay. Each strain was spotted 5-fold dilutions on glucose or galactose-containing plates to drive Hmg2-GFP and Dfm1-10xHis overexpression, and plates were incubated at 30°C.
E. Dilution assay as described in (A) except using rpn4Δ and ubp6Δ cells containing either GALpr-Hmg2-GFP, GALpr-Hmg2 (K6R)-GFP, GALpr-Hmg2 (K357R)-GFP, GALpr-Hmg2 (K6R and K357R)-GFP or EV.

**Figure S4: Chaperone Function of Dfm1 is Specific to Misfolded Proteins**

A. Western blot of aggregated versus soluble membrane proteins at the ER. Lysates from dfm1Δ cells containing SEC61-GFP, with either add-back of EV, WT DFM1, or DFM1-5Ashp were blotted with anti-GFP was to detect SEC61. One biological replicate was performed.

## METHODS

### Plasmids and Strains

Plasmids used in this study are listed in Table S1. Plasmids for this work were generated using standard molecular biological cloning techniques via polymerase chain reaction (PCR) of genes from yeast genomic DNA or plasmid followed by ligation into a specific restricted digested site within a construct and verified by sequencing (Eton Bioscience, Inc.). Primer information is available upon request.

A complete list of yeast strains and their corresponding genotypes are listed in Table S2. All strains used in this work were derived from S288C or Resgen. Yeast strains were transformed with DNA or PCR fragments using the standard LiOAc method in which null alleles were generated by using PCR to amplify a selection marker flanked by 30 base pairs of the 5’ and 3’ regions, which are immediately adjacent to the coding region of the gene to be deleted. The selectable markers used for making null alleles were genes encoding resistance to G418 or CloNat/nourseothricin or ability to synthesize histidine. After transformation, strains with drug markers were plated onto YPD followed by replica-plating onto YPD plates containing (500 μg/mL G418 or 200 μg/mL nourseothricin) or minimal media (-His) plates. All gene deletions were confirmed by PCR.

### Galactose Induction

For strains with plasmids containing galactose inducible promoters, protein expression was achieved by growing proteins overnight in appropriate selection media containing 2% raffinose as carbon source. The following day, samples were diluted between 0.10-0.20OD at 600nm (diluted absorbance was assay dependent). Cells in log phase were induced by adding 0.2% galactose to media. Minimum time requirement for robust protein expression was determined for strains using flow cytometry and was 2 or 3 hours for every strain used.

### Flow Cytometry

Yeast were grown in minimal medium with 2% raffinose and 0.2% galactose and appropriate amino acids into log phase (OD600 < 0.2). The BD Biosciences FACS Calibur flow cytometer measured the individual fluorescence of 10,000 cells. Experiments were analyzed using Prism8 (GraphPad).

### Unfolded Protein Response Activation Assay

Strains were inoculated overnight in minimal media (-His) with 2% raffinose. The following day, samples were diluted to 0.20OD in of minimal media (-His) and allowed to grow to log phase. Samples were then diluted to 0.30OD before adding 20% galactose to a final concentration of 0.2% galactose (+ GAL) or an equal volume of dH2O (-GAL). Timer was started after galactose addition and samples were measured using flow cytometry, as described above, every hour, starting from the 0-hour mark and ending at the 5-hour mark. At the 1-hour time point, samples were treated with either 2ug/mL tunicamycin or an equal volume of DMSO.

### Hac1 Splicing PCR

Strains were prepared the same as for the unfolded protein response activation assay, except they were grown in minimal media (-Ura -His) with 2% raffinose. After 5 hours of incubation with 0.2% galactose and 2ug/mL tunicamycin, samples were pelleted and washed with dH2O. RNA from samples was extracted using Qiagen RNeasy Mini Kit. Samples were ethanol precipitated by adding 1uL of glycoblue, 50uL of 7.5M ammonium acetate, and 700uL of chilled 100% ethanol. Tubes were then stored at −80°C for between three hours to overnight. Samples were then centrifuged at 13,000xg for 30 minutes at 4°C and supernatant was removed. Pellets were washed twice with 75% ethanol and centrifuged at room temperature at 13,000xg for 30 seconds. After drying the pellet, it was resuspended in 15uL of molecular grade water. 250ng of RNA from each sample was used to generate cDNA using a standard protocol for ProtoscriptII Reverse Transcriptase (NEB), except with 1uL of Oligo(dT)_12-18_ (Thermo Fisher Scientific) used for primer. Wizard SV Gel and PCR Clean-Up System (ProMega) was used on cDNA samples. Hac1 mRNA was amplified using forward primer 5’ACTTGGCTATCCCTACCAACT 3’ and reverse primer 5’ATGAATTCAAACCTGACTGC 3’. PCR products were resolved on a 2% agarose gel.

### MG132 Sensitivity Assay

MG132 sensitivity assay was performed using a protocol adapted from (Metzger, M.B. and Michaelis, S., 2009)^15^. In brief, cultures grown minimal media (-his) 2% dextrose. Cultures in log phase were split and treated with either 50uM MG132 in DMSO or an equal volume of DMSO alone and incubated for 8h at3 0°C. Cultures were diluted 1:500 and 100uL of sample was plated onto minimal media (-His) plates and grown at 30°C for 3 days. Experiment was done with two technical replicates and three biological replicates for each strain. Colony forming units (CFUs) were counted for DMSO-and MG132-treated cells using the ProMega Colony Counter application for iPhone.

### Spot dilution assay (Substrate-Toxicity Assay)

Yeast strains were grown in minimal selection media (-His) supplemented with 2% dextrose to log phase (OD600 0.2-0.3) at 30°C. 0.10 OD cells were pelleted and resuspended in 1mL dH2O. 250 μL of each sample was transferred to a 96-well plate where a five-fold serial dilution in dH2O of each sample was performed to obtain a gradient of 0.1-0.0000064 OD cells. The 8×6 pinning apparatus was used to pin cells onto synthetic complete (-His) agar plates supplemented with 2% dextrose or 2% galactose. Plates were incubated at 30°C and removed from the incubator for imaging after 3 days and again after 7 days.

### RNA Sequencing

RNA was isolated using a Qiagen RNeasy kit using standard protocol for yeast. Samples were eluted twice with 30uL of molecular grade water. To cleanup samples, 1uL of DNase was added to each sample and was incubated at 37°C for 25 minutes. 6uL of DNase inactivation buffer was added to samples and was incubated for 2 minutes. Samples were spun down at 10,000xg for 1.5 minutes and supernatant was transferred to a new microfuge tube. Samples were ethanol precipitated by adding 1uL of glycoblue, 50uL of 7.5M ammonium acetate, and 700uL of chilled 100% ethanol. Tubes were then stored at −80°C for between three hours to overnight. Samples were then centrifuged at 13,000xg for 30 minutes at 4°C and supernatant was removed. Pellets were washed twice with 75% ethanol and centrifuged at room temperature at 13,000xg for 30 seconds. After drying the pellet, it was resuspended in 15uL of molecular grade water.

Samples were measured for RNA concentration and an equal concentration of each sample was measured out into a total of 50uL of molecular grade water. RNA samples were added to washed, room temperature oligo dT beads in 50uL of buffer. Samples were poly-A enriched by incubating RNA with oligo dT beads at 65°C for 2 minutes and then incubating at room temperature 5-10 minutes. Beads were collected with magnet and washed twice with wash buffer 1 and wash buffer 2. 50uL of RNA elution buffer was added and then incubated at 80°C for 2 minutes. On ice, beads were collected on magnet and supernatant was removed and transferred to new chilled tubes. The beads were then washed with 200uL RNA elution buffer and 200uL 2x DTBB. Beads were resuspended in 50uL 2x DTBB and then RNA supernatant was transferred back to the tube with beads. Poly-A selection steps were repeated up until the wash with wash buffer 2 and samples were washed with 30uL of chilled 1x Superscript III first-strand buffer.

RNA was fragmented by resuspending beads in 10uL of fragmentation buffer and incubating at 94°C for 10 minutes. Beads were collected on magnet and supernatant was moved to new tubes. First strand cDNA synthesis was accomplished by adding 1.5uL of primer master mix and incubated at 55°C for 1 minute. 8.6uL of reverse transcriptase master mix was added to samples and then incubated at 25°C for 10 minutes followed by 50°C for 55 minutes. 36uL of RNAclean XP and isopropanol was added to samples and they were incubated on ice for 15 minutes. Beads were collected on magnet for 5 minutes and supernatant was discarded. Sample were washed twice with 80% ethanol and then RNA was eluted with 10uL of TET. To synthesize the 2^nd^ strand with dUTP, 5uL of 2^nd^ strand master mix was added to eluted RNA and incubated at 16°C for 2 hours to overnight. 5uL of dsDNA repair master mix was added and incubated at 20°C for 30 minutes. Samples were cleaned by added 30uL of speed beads in 20% PEG 8000 and 2.5M NaCl. Samples were spun down and 30uL isopropanol was added and incubated for 10-15 minutes. Beads were collected on magnets, supernatant was removed, and beads with washed twice with 80% ethanol and RNA was eluted in 16.1uL of warm TET. 13.9uL of dA tailing master mix was added to samples and they were incubated at 37°C for 30 minutes. 45uL of 20% PEG 8000 and 2.5M NaCl and an equal volume of isopropanol was added to samples and they were incubated at room temperature for 10-15 minutes. Beads were collected on magnets, supernatant was removed, and beads with washed twice with 80% ethanol and RNA was eluted in 13.7uL of warm TET

To add adapters, 15.8uL of ligation master mix was added to each sample on ice and 0.5uL of each unique barcode adapter was added to samples. Samples were incubated at room temperature for 15 minutes and were resuspended in 11uL of 20% PEG 8000 and 2.5M NaCl and they were incubated at room temperature for 10-15 minutes. Beads were collected on magnets, supernatant was removed, and beads with washed twice with 80% ethanol and RNA was eluted in 14uL of warm TET. UDG second strand digestion was accomplished by adding 1uL of UDG to samples and incubating at 37°C for 30 minutes.

Library was amplified by adding 10uL of PCR master mix to each sample and using the following PCR settings for five cycles: 98°C for 3 minutes (one time), 98°C for 45 seconds, 60°C for 30 seconds, and 72°C for 30 seconds, followed by 72°C for 3 minutes. Samples were cleaned by added 50uL of speed beads in 20% PEG 8000 and 2.5M NaCl. Beads were collected on magnets, supernatant was removed, and beads with washed twice with 80% ethanol and RNA was eluted in 15uL of warm TET.

To select library, samples were run on a 10% acrylamide 1xTBE gel. Gel was stained with cyber gold and bands were cut from 200-325bp. Samples were incubated overnight in gel elution buffer and samples were cleaned using Zymo columns according to manufacturer’s protocol, before eluting samples in 10uL of warm sequencing TET. Samples were sequenced using an Illumina HiSeq 2500.

### RNA Sequencing Data Analysis

Data was analyzed by normalizing reads per million and using principal components analysis to determine genes with the highest PC1 (+) scores and lowest PC1 (-) scores between *dfm1*Δ +GAL_pr_-Hmg2-GFP and every other strain tested. From this list, we used the top 100 genes with the highest (+) and lowest (-) PCA1 values, and cross referenced those to the normalized transcript per million reads value for each gene and removed genes that were not expressed at either a higher (for + PCA1 values) or lower (for - PCA1 values) reads per million level than all other conditions that were sequenced. Then, this list of upregulated and downregulated genes was used for gene ontology (GO) analysis using http://geneontology.org/. All code will be made available.

### Fluorescence Microscopy

To prepare cells, overnight cultures were diluted to ∼0.20 OD in minimal media lacking uracil (-URA). After growing ∼3 hours, samples were pelleted and washed with dH2O before being resuspended in 80uL of media to be used for imaging. Fluorescence microscopy was accomplished using a CSU-X1 Spinning Disk (Yokogawa) confocal microscope at the Nikon Imaging Center on the UCSD campus. Samples were analyzed to measure the fraction of GFP in puncta

### Microscopy Quantification and Analysis

Microscopy images were analyzed using ImageJ/Fiji (NIH). Images were thresholded and cells were manually traced in each image. Signals above the threshold were considered puncta. Mean gray value of all puncta in an individual cell were summed and divided by the mean gray value of the entire cell to determine fraction of Hmg2 in puncta for each cell. All statistical analysis was done using Prism8 (GraphPad). All code for analysis will be made available.

### Aggregation Assay

ER microsomes were isolated by centrifuging and pelleting 15OD of yeast in log phase growth. Pellets were resuspended in MF buffer with protease inhibitors and 0.5mM lysis beads were added to each sample. Samples were vortexed six times in 1-minute intervals, with 1-minute on ice in between. Lysed cells were transferred to new microcentrifuge tube and samples were clarified by spinning at 1,500x for 5 minutes at 4°C. Microsomes were separated by centrifuging clarified lysate at 14,000xg for 1 minute.. Fractions were incubated on ice in the presence or absence of 1% DDM for 1 hour. The mixture was then centrifuged at 16,000 × *g* for 30 min at 4°C, and the detergent soluble fraction (i.e., the supernatant) was precipitated with 20% TCA. Proteins from both the soluble and insoluble fractions were resuspended in sample buffer and resolved by SDS-PAGE.

### Western Blot Quantification

Western blot images were quantified using ImageJ/Fiji. Band intensities were measured from high resolution TIF files of western blot images acquired from a BioRad Chemidoc Imager. Statistical analysis was done using Prism8 (GraphPad).

