## Supplementary figures and images for "Derlin Dfm1 Employs a Chaperone Function to Resolve Misfolded Membrane Protein Stress"

### Supplemental Figure 1

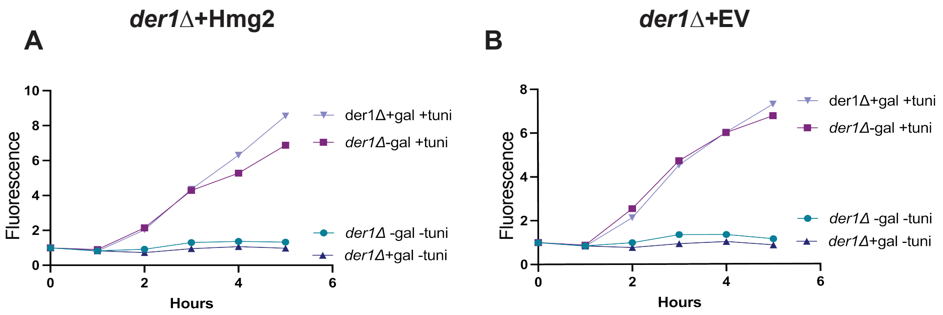


Supplemental Figure 1, Kandel et al.

### Supplemental Figure 2

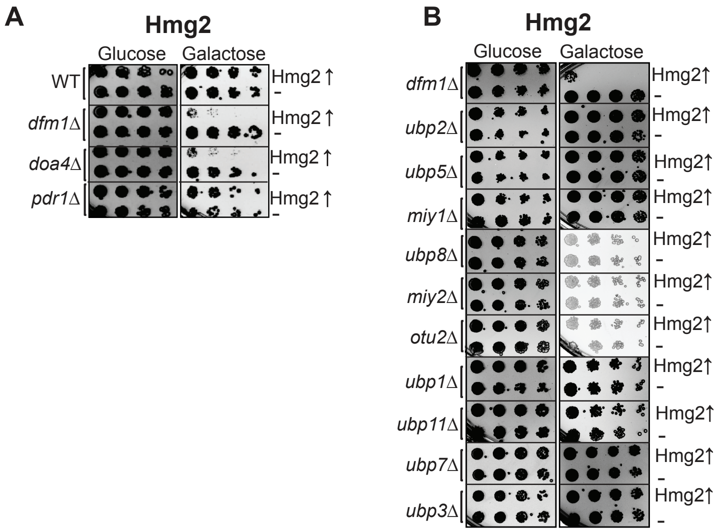


Supplemental Figure 2, Kandel et al.

### Supplemental Figure 3

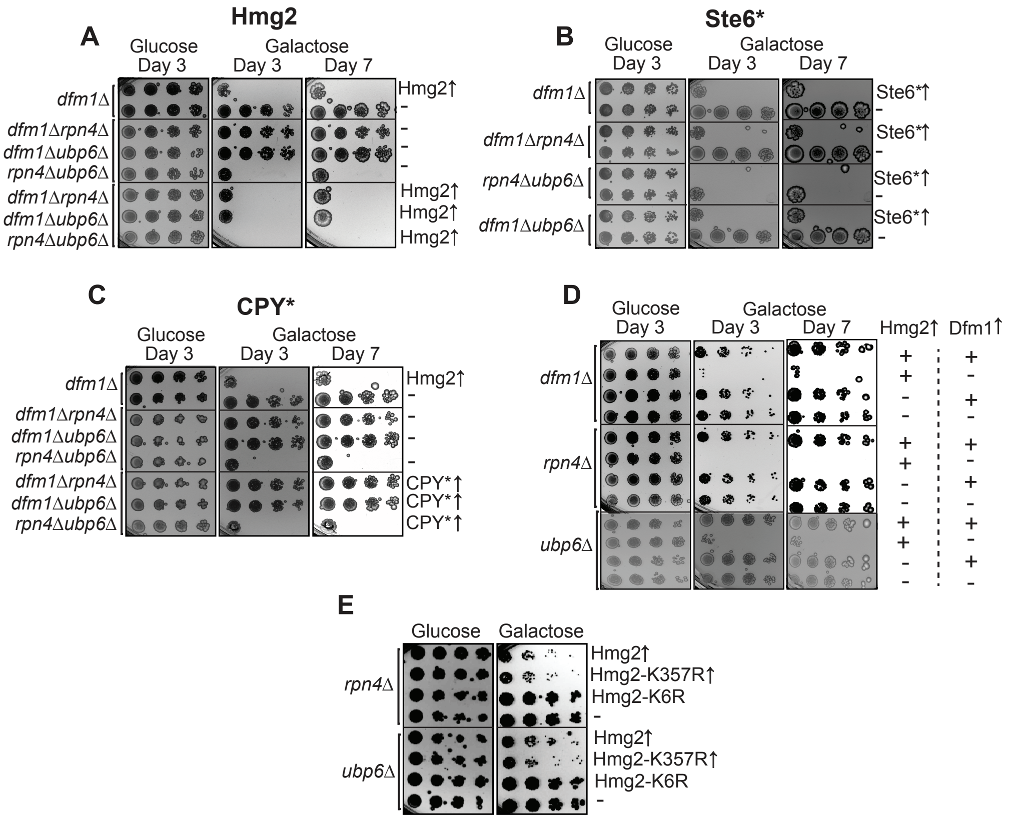


Supplemental Figure 3, Kandel et al.

### Supplemental Figure 4

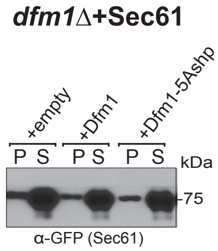


Supplemental Figure 4, Kandel et al.
