## Supplemental Table 1 for "Derlin Dfm1 Employs a Chaperone Function to Resolve Misfolded Membrane Protein Stress"

**Table S1. Plasmids used in this study, Related to Figures 1-8**

| Plasmid | Gene | | |
|---|---|---|---|
| pRH1120 | YCp | URA3 | pGAL1-HMG2-GFP |
| pRH3113 | YCp | URA3 | pGAL1-PDR5*-HA |
| pRH3112 | YCp | URA3 | pGAL1-CPY*-HA |
| pRH3144 | YCp | URA3 | pGAL1- Ste6-166p-3HA-GFP |
| pRH317 | YCp | URA3 | |
| pRH316 | YCp | LEU2 | |
| pRH1945 | YIp | ADE2 URA3 | 4xUPRE-GFP |
| pRH2890 | YCp | LEU2 | pDFM1-DFM1-3HA-5aShp |
| pRH1997 | YCp | LEU2 | pDFM1-DFM1-3HA |
| pRH2812 | YCp | LEU2 | pDFM1-DFM1-3HA-AxxxG |
| pRH2813 | YCp | LEU2 | pDFM1-DFM1-3HA-GxxxA |
| pRH2826 | YCp | LEU2 | pDFM1-DFM1-3HA-WA |
| pRH2827 | YCp | LEU2 | pDFM1-DFM1-3HA-AR |
| pRH2013 | YCp | LEU2 | pDFM1-DFM1-3HA |
| pSN12 | YIp | ADE2 HIS3 | |
| pSN11 | YCp | URA3 | |
| pSN103 | YIp | ADE2 HIS3 | pGAL1-HMG2-GFP-K357R |
| pSN104 | YIp | ADE2 HIS3 | pGAL1-HMG2-GFP-K6R |

| | | | |
|---|---|---|---|
| pSN105 | YIp | ADE2 HIS3 | pGAL1-HMG2-GFP |
| pSN59 | YCp | LEU2 | pDFM1-DFM1-3HA-F107S |
| pSN60 | YCp | LEU2 | pDFM1-DFM1-3HA-L64V |
| pSN93 | YCp | LEU2 | pDFM1-DFM1-3HA-K67E |
| pSN94 | YCp | LEU2 | pDFM1-DFM1-3HA-Q101R |
| pSN95 | YCp | LEU2 | pDFM1-DFM1-3HA-F57S |
| pSN168 | YCp | ADE2 HIS3 | pGAL1-HMG2-GFP-K6R-K357R |
| pSN86 | YIp | ADE2 HIS3 | pGAL1-STE6-166p-3HA-GFP |
| pSN88 | YIp | ADE2 HIS3 | pGAL1-HMG2-6MYC |
| pSN100 | YIp | ADE2 HIS3 | pGAL1-CPY*-HA |
| pSN195 | YCp | URA3 | CFTR-HA |
| pSN196 | YCp | URA3 | CFTR-HA-ΔF508 |
| pSN197 | YCp | URA3 | A1PiZ |
| pSN199 | YCp | URA3 | pGAL1-DFM1-6HIS |
| pSN5 | YCp | URA3 | |
| pSN193 | YIp | ADE2 LEU2 | pADH1-Derlin-1-MYC |
| pSN194 | YIp | ADE2 LEU2 | pADH1-Derlin-2-MYC |
| pSN39 | YIp | ADE2 LEU2 | |
| pSN190 | YCp | URA3 | pCUP1-HBT-Ubiquitin |
| pRH613 | YIp | ADE2 | pTDH3-Hmg2-GFP |

| | | | |
|---|---|---|---|
| pRH2058 | YCp | URA3 | pPGK1-STE6-166p-3HA-GFP |
| pRH311 | YIp | TRP1 | |
