## Supplemental Table 2 for "Derlin Dfm1 Employs a Chaperone Function to Resolve Misfolded Membrane Protein Stress"

**Table S2. Yeast strains used in this study, Related to Figures 1-6**

| Strain | Genotype | Reference |
|---|---|---|
| RHY10520 | Mata *ADE2 met15Δ0 LYS2(LYS+) ura3Δ0 TRP1 leu2Δ0 his3Δ1 pdr5Δ::KanMX CEN::URA3* | This study |
| RHY10519 | *Mata ADE2 met15Δ0 LYS2(LYS+) ura3Δ0 TRP1 leu2Δ0 his3Δ1 pdr5Δ::KanMX CEN::URA3::GAL1pr-HMG2-GFP* | This study |
| RHY10518 | *Mata ADE2 met15Δ0 LYS2(LYS+) ura3Δ0 TRP1 leu2Δ0 his3Δ1 dfm1Δ::KanMX CEN::URA3* | This study |
| RHY10517 | *Mata ADE2 met15Δ0 LYS2(LYS+) ura3Δ0 TRP1 leu2Δ0 his3Δ1 dfm1Δ::KanMX CEN::URA3::GAL1pr-HMG2-GFP* | This study |
| RHY10655 | *Mata ADE2 met15Δ0 LYS2(LYS+) ura3Δ0 TRP1 leu2Δ0 his3Δ1hrd1Δ::KanMX CEN::URA3* | This study |
| RHY10654 | *Mata ADE2 met15Δ0 LYS2(LYS+) ura3Δ0 TRP1 leu2Δ0 his3Δ1 hrd1Δ::KanMX CEN::URA3::GAL1pr-HMG2-GFP* | This study |
| RHY11580 | *Mata ADE2 met15Δ0 LYS2(LYS+) ura3Δ0 TRP1 leu2Δ0 his3Δ1 pdr5Δ::KanMX CEN::URA3::GAL1pr-PDR5*-HA* | This study |
| RHY11581 | *Mata ADE2 met15Δ0 LYS2(LYS+) ura3Δ0 TRP1 leu2Δ0 his3Δ1 dfm1Δ::KanMX CEN::URA3::GAL1pr-PDR5*-HA* | This study |
| RHY11583 | *Mata ADE2 met15Δ0 LYS2(LYS+) ura3Δ0 TRP1 leu2Δ0 his3Δ1 hrd1Δ::KanMX CEN::URA3::GAL1pr-PDR5*-HA* | This study |
| RHY11576 | *Mata ADE2 met15Δ0 LYS2(LYS+) ura3Δ0 TRP1 leu2Δ0 his3Δ1 pdr5Δ::KanMX CEN::URA3::GAL1pr-CPY*-HA* | This study |
| RHY11577 | *Mata ADE2 met15Δ0 LYS2(LYS+) ura3Δ0 TRP1 leu2Δ0 his3Δ1 dfm1Δ::KanMX CEN::URA3::GAL1pr-CPY*-HA* | Neal et al., 2020 |
| RHY11579 | *Mata ADE2 met15Δ0 LYS2(LYS+) ura3Δ0 TRP1 leu2Δ0 his3Δ1 hrd1Δ::KanMX CEN::URA3::GAL1pr-CPY*-HA* | Neal et al., 2020 |
| RHY11817 | *Mata ADE2 MET2 lys2-801 ura3Δ0 TRP1 leu2Δ0 his3Δ200 dfm1Δ::KanMX hrd1Δ::CloNAT CEN::URA3* | Neal et al., 2020 |
| RHY11818 | *Mata ADE2 MET2 lys2-801 ura3Δ0 TRP1 leu2Δ0 his3Δ200 dfm1Δ::KanMX hrd1Δ::CloNAT CEN::URA3::GAL1pr-HMG2-GFP* | This study |

| | | |
|---|---|---|
| RHY 11867 | *Mata ADE2 met15Δ0 LYS2(LYS+) ura3Δ0 TRP1 leu2Δ0 his3Δ1 pdr5Δ::KanMX*<br>*CEN::URA3::GAL1pr-STE6-166p-3HA-GFP* | Neal et al., 2018 |
| RHY 11868 | *Mata ADE2 met15Δ0 LYS2(LYS+) ura3Δ0 TRP1 leu2Δ0 his3Δ1 dfm1Δ::KanMX*<br>*CEN::URA3::GAL1pr-STE6-166p-3HA-GFP* | Neal et al., 2018 |
| RHY 11869 | *Mata ADE2 met15Δ0 LYS2(LYS+) ura3Δ0 TRP1 leu2Δ0 his3Δ1 hrd1Δ::KanMX*<br>*CEN::URA3::GAL1pr-STE6-166p-3HA-GFP* | Neal et al., 2020 |
| RHY 11826 | *Mata ADE2 MET2 lys2-801 ura3Δ0 TRP1 leu2Δ0 his3Δ200 dfm1Δ::KanMX hrd1Δ::CloNAT*<br>*CEN::URA3::GAL1pr-STE6-166p-3HA-GFP* | This study |
| RHY 11873 | *Mata ADE2 met15Δ0 LYS2(LYS+) ura3-52 trp1::hisG leu2Δ his3Δ1 dfm1Δ::CloNAT doa10Δ::HphMx*<br>*CEN::URA3::GAL1pr-STE6-166p-3HA-GFP* | This study |
| RHY 11908 | *Mata ADE2 met15Δ0 LYS2(LYS+) ura3-52 trp1::hisG leu2Δ his3Δ1 doa10Δ::HphMx*<br>*CEN::URA3::GAL1pr-STE6-166p-3HA-GFP* | This study |
| RHY 11906 | *Mata ADE2 met15Δ0 LYS2(LYS+) ura3-52 trp1::hisG leu2Δ his3Δ1 doa10Δ::HphMx*<br>*CEN::URA3* | Neal et al., 2020 |
| RHY 11907 | *Mata ADE2 met15Δ0 LYS2(LYS+) ura3-52 trp1::hisG leu2Δ his3Δ1 doa10Δ::HphMx*<br>*CEN::URA3::GAL1pr-HMG2-GFP* | Neal et al., 2020 |
| RHY 11914 | *Mata ADE2 met15Δ0 LYS2(LYS+) ura3-52 trp1::hisG leu2Δ his3Δ1 dfm1Δ::CloNAT doa10Δ::HphMx*<br>*CEN::URA3* | This study |
| RHY 11915 | *Mata ADE2 met15Δ0 LYS2(LYS+) ura3-52 trp1::hisG leu2Δ his3Δ1 dfm1Δ::CloNAT doa10Δ::HphMx*<br>*CEN::URA3::GAL1pr-HMG2-GFP* | This study |
| RHY 11226 | *Mata ADE2 met15Δ0 LYS2(LYS+) ura3Δ0 TRP1 leu2Δ0 his3Δ1 dfm1Δ::KanMX*<br>*CEN::URA3 CEN::LEU2::pDFM1-DFM1-3HA* | Neal et al., 2018 |
| RHY 11227 | *Mata ADE2 met15Δ0 LYS2(LYS+) ura3Δ0 TRP1 leu2Δ0 his3Δ1 dfm1Δ::KanMX*<br>*CEN::URA3:: GAL1pr-HMG2-GFP CEN::LEU2::pDFM1-DFM1-3HA* | Neal et al., 2018 |
| RHY 11216 | *Mata ADE2 met15Δ0 LYS2(LYS+) ura3Δ0 TRP1 leu2Δ0 his3Δ1 dfm1Δ::KanMX*<br>*CEN::URA3 CEN::LEU2* | Neal et al., 2018 |
| RHY11217 | *Mata ADE2 met15Δ0 LYS2(LYS+) ura3Δ0 TRP1 leu2Δ0 his3Δ1 dfm1Δ::KanMX*<br>*CEN::URA3:: GAL1pr-HMG2-GFP CEN::LEU2* | Neal et al., 2018 |

| | | |
|---|---|---|
| RHY11222 | *Mata ADE2 met15Δ0 LYS2(LYS+) ura3Δ0 TRP1 leu2Δ0 his3Δ1 dfm1Δ::KanMX CEN::URA3 CEN::LEU2::pDFM1-DFM1-3HA-WA* | Neal et al., 2018 |
| RHY11223 | *Mata ADE2 met15Δ0 LYS2(LYS+) ura3Δ0 TRP1 leu2Δ0 his3Δ1 dfm1Δ::KanMX CEN::URA3:: GAL1pr-HMG2-GFP CEN::LEU2::pDFM1-DFM1-3HA-WA* | Neal et al., 2018 |
| RHY11224 | *Mata ADE2 met15Δ0 LYS2(LYS+) ura3Δ0 TRP1 leu2Δ0 his3Δ1 dfm1Δ::KanMX CEN::URA3 CEN::LEU2::pDFM1-DFM1-3HA-AR* | Neal et al., 2018 |
| RHY11225 | *Mata ADE2 met15Δ0 LYS2(LYS+) ura3Δ0 TRP1 leu2Δ0 his3Δ1 dfm1Δ::KanMX CEN::URA3:: GAL1pr-HMG2-GFP CEN::LEU2::pDFM1-DFM1-3HA-AR* | Neal et al., 2018 |
| RHY11218 | *Mata ADE2 met15Δ0 LYS2(LYS+) ura3Δ0 TRP1 leu2Δ0 his3Δ1 dfm1Δ::KanMX CEN::URA3 CEN::LEU2::pDFM1-DFM1-3HA-AxxxG* | Neal et al., 2018 |
| RHY11219 | *Mata ADE2 met15Δ0 LYS2(LYS+) ura3Δ0 TRP1 leu2Δ0 his3Δ1 dfm1Δ::KanMX CEN::URA3:: GAL1pr-HMG2-GFP CEN::LEU2::pDFM1-DFM1-3HA-AxxxG* | Neal et al., 2018 |
| RHY11220 | *Mata ADE2 met15Δ0 LYS2(LYS+) ura3Δ0 TRP1 leu2Δ0 his3Δ1 dfm1Δ::KanMX CEN::URA3 CEN::LEU2::pDFM1-DFM1-3HA-GxxxA* | Neal et al., 2018 |
| RHY11221 | *Mata ADE2 met15Δ0 LYS2(LYS+) ura3Δ0 TRP1 leu2Δ0 his3Δ1 dfm1Δ::KanMX CEN::URA3:: GAL1pr-HMG2-GFP CEN::LEU2::pDFM1-DFM1-3HA-GxxxA* | Neal et al., 2018 |
| RHY11073 | *Mata ADE2 met15Δ0 LYS2(LYS+) ura3Δ0 TRP1 leu2Δ0 his3Δ1 dfm1Δ::KanMX CEN::URA3 CEN::LEU2::pDFM1-DFM1-3HA-5aShp* | Neal et al., 2018 |
| RHY11074 | *Mata ADE2 met15Δ0 LYS2(LYS+) ura3Δ0 TRP1 leu2Δ0 his3Δ1 dfm1Δ::KanMX CEN::URA3:: GAL1pr-HMG2-GFP CEN::LEU2::pDFM1-DFM1-3HA-5aShp* | Neal et al., 2018 |
| SEN141 | Mata *ade2::ADE2::HIS3::pGAL1::Hmg2-GFP met2 lys2-801 ura3-52 trp1::hisG leu2Δ his3Δ200* | This study |
| SEN142 | Mata *ade2::ADE2::HIS3 met2 lys2-801 ura3-5, trp1::hisG leu2Δ his3Δ200* | This study |
| SEN149 | Mata *ade2::ADE2::HIS3 met2 lys2-801 ura3-52 trp1::hisG leu2Δ his3Δ200 dfm1Δ::KanMX* | This study |
| SEN165 | Mata *ade2::ADE2::HIS3::pGAL1::Hmg2-GFP met2 lys2-801 ura3-52, trp1::hisG leu2Δ his3Δ200 dfm1Δ::KanMX* | This study |
| SEN407 | Mata *ade2-101 met2 lys2-801 ura3-5, trp1::hisG leu2Δ his3Δ200* | This study |

| | | |
|---|---|---|
| | *CEN::ADE2::HIS3::pGAL1-Hmg2-GFP-K6R-K357R* | |
| SEN408 | Mata *ade2-101 met2 lys2-801 ura3-5, trp1::hisG leu2Δ his3Δ200 dfm1Δ::KanMX CEN::ADE2::HIS3::pGAL1-Hmg2-GFP-K6R-K357R* | This study |
| SEN139 | Mata *ade2::ADE2::HIS3::pGAL1::Hmg2-GFP-K357R met2 lys2-801 ura3-52 trp1::hisG leu2Δ his3Δ200* | This study |
| SEN147 | Mata *ade2::ADE2::HIS3::pGAL1::Hmg2-GFP-K357R met2 lys2-801 ura3-52 trp1::hisG leu2Δ his3Δ200 dfm1Δ::KanMX* | This study |
| SEN140 | Mata *ade2::ADE2::HIS3::pGAL1::Hmg2-GFP-K6R met2 lys2-801 ura3-52 trp1::hisG leu2Δ his3Δ200* | This study |
| SEN148 | Mata *ade2::ADE2::HIS3::pGAL1::Hmg2-GFP-K6R met2 lys2-801 ura3-52 trp1::hisG leu2Δ his3Δ200 dfm1Δ::KanMX* | This study |
| SEN182 | Mata *ade2::ADE2::HIS3 met2 lys2-801 ura3-52 trp1::hisG leu2Δ his3Δ200 dfm1Δ::KanMX CEN::LEU2::pDFM1-DFM1-3HA* | This study |
| SEN183 | Mata *ade2::ADE2::HIS3 met2 lys2-801 ura3-52 trp1::hisG leu2Δ his3Δ200 dfm1Δ::KanMX CEN::LEU2* | This study |
| SEN192 | Mata *ade2::ADE2::HIS3::pGAL1::Hmg2-GFP met2 lys2-801 ura3-52, trp1::hisG leu2Δ his3Δ200 dfm1Δ::KanMX CEN::LEU2::pDFM1-DFM1-3HA* | This study |
| SEN193 | Mata *ade2::ADE2::HIS3::pGAL1::Hmg2-GFP met2 lys2-801 ura3-52, trp1::hisG leu2Δ his3Δ200 dfm1Δ::KanMX CEN::LEU2* | This study |
| SEN250 | Mata *ade2::ADE2::HIS3 met2 lys2-801 ura3-52 trp1::hisG leu2Δ his3Δ200 dfm1Δ::KanMX CEN::LEU2::pDFM1-DFM1-3HA-F107S* | This study |
| SEN251 | Mata *ade2::ADE2::HIS3 met2 lys2-801 ura3-52 trp1::hisG leu2Δ his3Δ200 dfm1Δ::KanMX CEN::LEU2::pDFM1-DFM1-3HA-L64V* | This study |
| SEN252 | Mata *ade2::ADE2::HIS3 met2 lys2-801 ura3-52 trp1::hisG leu2Δ his3Δ200 dfm1Δ::KanMX CEN::LEU2::pDFM1-DFM1-3HA-K67E* | This study |
| SEN253 | Mata *ade2::ADE2::HIS3 met2 lys2-801 ura3-52 trp1::hisG leu2Δ his3Δ200 dfm1Δ::KanMX CEN::LEU2::pDFM1-DFM1-3HA-Q101R* | This study |
| SEN254 | Mata *ade2::ADE2::HIS3 met2 lys2-801 ura3-52 trp1::hisG leu2Δ his3Δ200 dfm1Δ::KanMX CEN::LEU2::pDFM1-DFM1-3HA-F58S* | This study |
| SEN256 | Mata *ade2::ADE2::HIS3::pGAL1::Hmg2-GFP met2 lys2-801 ura3-52, trp1::hisG leu2Δ his3Δ200 dfm1Δ::KanMX CEN::LEU2::pDFM1-DFM1-3HA-F107S* | This study |
| SEN257 | Mata *ade2::ADE2::HIS3::pGAL1::Hmg2-GFP met2 lys2-801 ura3-52, trp1::hisG leu2Δ his3Δ200 dfm1Δ::KanMX* | This study |

| | | |
|---|---|---|
| | *CEN::LEU2::pDFM1-DFM1-3HA-L64V* | |
| SEN258 | Mata *ade2::ADE2::HIS3::pGAL1::Hmg2-GFP met2 lys2-801 ura3-52, trp1::hisG leu2Δ his3Δ200 dfm1Δ::KanMX CEN::LEU2::pDFM1-DFM1-3HA-K67E* | This study |
| SEN259 | Mata *ade2::ADE2::HIS3::pGAL1::Hmg2-GFP met2 lys2-801 ura3-52, trp1::hisG leu2Δ his3Δ200 dfm1Δ::KanMX CEN::LEU2::pDFM1-DFM1-3HA-Q101R* | This study |
| SEN260 | Mata *ade2::ADE2::HIS3::pGAL1::Hmg2-GFP met2 lys2-801 ura3-52, trp1::hisG leu2Δ his3Δ200 dfm1Δ::KanMX CEN::LEU2::pDFM1-DFM1-3HA-F58S* | This study |
| SEN103 | Mata *ade2::ADE2::URA3::4xUPRE-GFP met15Δ0 LYS2 (LYS+) ura3Δ0 TRP1 leu2Δ0 his3Δ1 pdr5Δ::KanMX pGAL1::CPY*-HA* | This study |
| SEN111 | Mata *ade2::ADE2::URA3::4xUPRE-GFP met15Δ0 LYS2 (LYS+) ura3Δ0 TRP1 leu2Δ0 his3Δ1 dfm1Δ:: KanMX pGAL1::CPY*-HA* | This study |
| SEN73 | Mata *ade2::ADE2::URA3::4xUPRE-GFP met15Δ0 LYS2 (LYS+) ura3Δ0 TRP1 leu2Δ0 his3Δ1 pdr5Δ::KanMX pGAL1::STE6-166p-3HA-GFP* | This study |
| SEN75 | Mata *ade2::ADE2::URA3::4xUPRE-GFP met15Δ0 LYS2 (LYS+) ura3Δ0 TRP1 leu2Δ0 his3Δ1 dfm1Δ:: KanMX pGAL1::STE6-166p-3HA-GFP* | This study |
| SEN76 | Mata *ade2::ADE2::URA3::4xUPRE-GFP met15Δ0 LYS2 (LYS+) ura3Δ0 TRP1 leu2Δ0 his3Δ1 ADE2::HIS3 pdr5Δ::KanMX* | This study |
| SEN68 | Mata *ade2::ADE2::URA3::4xUPRE-GFP met15Δ0 LYS2 (LYS+) ura3Δ0 TRP1 leu2Δ0 his3Δ1 ADE2::HIS3 dfm1Δ::KanMX* | This study |
| SEN70 | Mata *ade2::ADE2::URA3::4xUPRE-GFP met15Δ0 LYS2 (LYS+) ura3Δ0 TRP1 leu2Δ0 his3Δ1 dfm1Δ::KanMX pGAL1::HMG2::6MYC* | This study |
| SEN71 | Mata *ade2::ADE2::URA3::4xUPRE-GFP met15Δ0 LYS2 (LYS+) ura3Δ0 TRP1 leu2Δ0 his3Δ1 ADE2::HIS3 dfm1Δ::KanMX* | This study |
| SEN155 | Mata *ADE2::HIS3 met15Δ0 LYS2 (LYS+) ura3Δ0 TRP1 leu2Δ0 his3Δ1 rpn4Δ::KanMX* | This study |
| SEN166 | Mata *ADE2::HIS3::pGAL::Hmg2-GFP met15Δ0 LYS2 (LYS+) ura3Δ0 TRP1 leu2Δ0 his3Δ1 rpn4Δ::KanMX* | This study |
| SEN196 | Mata *ADE2::HIS3 met15Δ0 LYS2 (LYS+) ura3Δ0 TRP1 leu2Δ0 his3Δ1 ubp6Δ::KanMX* | This study |
| SEN197 | Mata *ADE2::HIS3::pGAL1::Hmg2-GFP met15Δ0 LYS2 (LYS+) ura3Δ0 TRP1 leu2Δ0 his3Δ1 ubp6Δ::KanMX* | This study |
| SEN411 | Mata *ade2::ADE2::HIS3::pGAL1::STE6-166-3HA-GFP met2 lys2-801 ura3-52 trp1::hisG leu2Δ his3Δ200* | This study |

| | | |
|---|---|---|
| SEN412 | Mata ADE2::HIS3::pGAL1::STE6-166p-3HA-GFP met15Δ0 LYS2 (LYS+) ura3Δ0 TRP1 leu2Δ0 his3Δ1 rpn4Δ::KanMX | This study |
| SEN413 | Mata ade2::ADE2::HIS3:: pGAL1::STE6-166p-3HA-GFP met2 lys2-801 ura3-52 trp1::hisG leu2Δ his3Δ200 dfm1Δ::KanMX | This study |
| SEN414 | Mata ADE2::HIS3::pGAL1::STE6-166p-3HA-GFP met15Δ0 LYS2 (LYS+) ura3Δ0 TRP1 leu2Δ0 his3Δ1 ubp6Δ::KanMX | This study |
| SEN269 | Mata ADE2::HIS3::pGAL1::CPY*-HA met15Δ0 LYS2 (LYS+) ura3Δ0 TRP1 leu2Δ0 his3Δ1 ubp6Δ::KanMX | This study |
| SEN415 | Mata ade2::ADE2::HIS3::pGAL1::CPY*-HA met2 lys2-801 ura3-52 trp1::hisG leu2Δ his3Δ200 | This study |
| SEN416 | Mata ADE2::HIS3::pGAL1::CPY*-HA met15Δ0 LYS2 (LYS+) ura3Δ0 TRP1 leu2Δ0 his3Δ1 rpn4Δ::KanMX | This study |
| SEN417 | Mata ade2::ADE2::HIS3:: pGAL1::CPY*-HA met2 lys2-801 ura3-52 trp1::hisG leu2Δ his3Δ200 dfm1Δ::KanMX | This study |
| SEN270 | Mata ADE2::HIS3::pGAL1::CPY*-HA met15Δ0 LYS2 (LYS+) ura3Δ0 TRP1 leu2Δ0 his3Δ1 rpn4Δ::KanMX dfm1Δ::CloNAT | This study |
| SEN271 | Mata ADE2::HIS3::pGAL1::CPY*-HA met15Δ0 LYS2 (LYS+) ura3Δ0 TRP1 leu2Δ0 his3Δ1 rpn4Δ::KanMX ubp6Δ::CloNAT | This study |
| SEN272 | Mata ADE2::HIS3::pGAL1::CPY*-HA met15Δ0 LYS2 (LYS+) ura3Δ0 TRP1 leu2Δ0 his3Δ1 ubp6Δ::KanMX dfm1Δ::CloNAT | This study |
| SEN499 | Mata ADE2::HIS3::pGAL1::STE6-166p-3HA-GFP met15Δ0 LYS2 (LYS+) ura3Δ0 TRP1 leu2Δ0 his3Δ1 rpn4Δ::KanMX dfm1Δ::CloNAT | This study |
| SEN500 | Mata ADE2::HIS3::pGAL1::STE6-166p-3HA-GFP met15Δ0 LYS2 (LYS+) ura3Δ0 TRP1 leu2Δ0 his3Δ1 rpn4Δ::KanMX ubp6Δ::CloNAT | This study |
| SEN501 | Mata ADE2::HIS3::pGAL1::STE6-166p-3HA-GFP met15Δ0 LYS2 (LYS+) ura3Δ0 TRP1 leu2Δ0 his3Δ1 ubp6Δ::KanMX dfm1Δ::CloNAT | This study |
| SEN273 | Mata ADE2::HIS3::pGAL1::HMG2-GFP met15Δ0 LYS2 (LYS+) ura3Δ0 TRP1 leu2Δ0 his3Δ1 rpn4Δ::KanMX dfm1Δ::CloNAT | This study |
| SEN276 | Mata ADE2::HIS3 met15Δ0 LYS2 (LYS+) ura3Δ0 TRP1 leu2Δ0 his3Δ1 rpn4Δ::KanMX dfm1Δ::CloNAT | This study |
| SEN274 | Mata ADE2::HIS3::pGAL1::HMG2-GFP met15Δ0 LYS2 (LYS+) ura3Δ0 TRP1 leu2Δ0 his3Δ1 rpn4Δ::KanMX ubp6Δ::CloNAT | This study |
| SEN277 | Mata ADE2::HIS3 met15Δ0 LYS2 (LYS+) ura3Δ0 TRP1 leu2Δ0 his3Δ1 rpn4Δ::KanMX ubp6Δ::CloNAT | This study |

| | | |
|---|---|---|
| SEN275 | Mata *ADE2::HIS3::pGAL1::HMG2-GFP met15Δ0 LYS2 (LYS+) ura3Δ0 TRP1 leu2Δ0 his3Δ1 ubp6Δ::KanMX dfm1Δ::CloNAT* | This study |
| SEN278 | Mata *ADE2::HIS3 met15Δ0 LYS2 (LYS+) ura3Δ0 TRP1 leu2Δ0 his3Δ1 ubp6Δ::KanMX dfm1Δ::CloNAT* | This study |
| SEN487 | Mata *ade2::ADE2::HIS3::pGAL1::Hmg2-GFP met2 lys2-801 ura3-52, trp1::hisG leu2Δ his3Δ200 dfm1Δ::KanMX CEN::URA3::pGAL1::DFM1-6HIS* | This study |
| SEN488 | Mata *ade2::ADE2::HIS3::pGAL1::Hmg2-GFP met2 lys2-801 ura3-52, trp1::hisG leu2Δ his3Δ200 dfm1Δ::KanMX CEN::URA3* | This study |
| SEN489 | Mata *ade2::ADE2::HIS3 met2 lys2-801 ura3-52, trp1::hisG leu2Δ his3Δ200 dfm1Δ::KanMX CEN::URA3::pGAL1::DFM1-6HIS* | This study |
| SEN490 | Mata *ade2::ADE2::HIS3 met2 lys2-801 ura3-52, trp1::hisG leu2Δ his3Δ200 dfm1Δ::KanMX CEN::URA3* | This study |
| SEN491 | Mata *ADE2::HIS3::pGAL::Hmg2-GFP met15Δ0 LYS2 (LYS+) ura3Δ0 TRP1 leu2Δ0 his3Δ1 rpn4Δ::KanMX CEN::URA3::pGAL1::DFM1-6HIS* | This study |
| SEN492 | Mata *ADE2::HIS3::pGAL::Hmg2-GFP met15Δ0 LYS2 (LYS+) ura3Δ0 TRP1 leu2Δ0 his3Δ1 rpn4Δ::KanMX CEN::URA3* | This study |
| SEN493 | Mata *ADE2::HIS3 met15Δ0 LYS2 (LYS+) ura3Δ0 TRP1 leu2Δ0 his3Δ1 rpn4Δ::KanMX CEN::URA3::pGAL1::DFM1-6HIS* | This study |
| SEN494 | Mata *ADE2::HIS3 met15Δ0 LYS2 (LYS+) ura3Δ0 TRP1 leu2Δ0 his3Δ1 rpn4Δ::KanMX CEN::URA3* | This study |
| SEN517 | Mata *ADE2::HIS3::pGAL::Hmg2-GFP met15Δ0 LYS2 (LYS+) ura3Δ0 TRP1 leu2Δ0 his3Δ1 ubp6Δ::KanMX CEN::URA3::pGAL1::DFM1-6HIS* | This study |
| SEN518 | Mata *ADE2::HIS3::pGAL::Hmg2-GFP met15Δ0 LYS2 (LYS+) ura3Δ0 TRP1 leu2Δ0 his3Δ1 ubp6Δ::KanMX CEN::URA3* | This study |
| SEN519 | Mata *ADE2::HIS3 met15Δ0 LYS2 (LYS+) ura3Δ0 TRP1 leu2Δ0 his3Δ1 ubp6Δ::KanMX CEN::URA3::pGAL1::DFM1-6HIS* | This study |
| SEN520 | Mata *ADE2::HIS3 met15Δ0 LYS2 (LYS+) ura3Δ0 TRP1 leu2Δ0 his3Δ1 ubp6Δ::KanMX CEN::URA3* | This study |
| SEN249 | Mata *ade2-101 met2 lys2-801 ura3-52 trp1::hisG::TRP1:: pTDH3-Hmg1p-myc-Hrd1p-3HA-GFP leu2Δ his3Δ200 hrd1Δ::KanMX dfm1Δ::CloNAT pdr5Δ::HIS3* | This study |
| SEN378 | Mata *ade2-101 met2 lys2-801 ura3-52 trp1::hisG::TRP1 leu2Δ his3Δ200 hrd1Δ::KanMX pdr5Δ::HIS3* | This study |

| | | |
|---|---|---|
| SEN229 | Mata *ade2-101 met2 lys2-801 ura3-52 trp1::hisG::TRP1::pTDH3-Hmg1p-myc-Hrd1p-3HA-GFP leu2Δ his3Δ200 hrd1Δ::KanMX pdr5Δ::HIS3* | This study |
| SEN228 | Mata *ade2-101 met2 lys2-801 ura3-52 trp1::hisG::TRP1::pTDH3-Hmg1p-myc-Hrd1p-3HA-GFP leu2Δ his3Δ200 pdr5Δ::HIS3* | This study |
| SEN377 | Mata *ade2-101 met2 lys2-801 ura3-52 trp1::hisG::TRP1 leu2Δ his3Δ200 pdr5Δ::HIS3* | This study |
| SEN379 | Mata *ade2-101 met2 lys2-801 ura3-52 trp1::hisG::TRP1 leu2Δ his3Δ200 hrd1Δ::KanMX dfm1Δ::CloNAT pdr5Δ::HIS3* | This study |
| SEN401 | Mata *ADE2::HIS3::pGAL1::Hmg2-GFP met15Δ0 LYS2 (LYS+) ura3Δ0 TRP1 leu2Δ0 his3Δ1 ubp9Δ::KanMX* | This study |
| SEN424 | Mata *ADE2::HIS3 met15Δ0 LYS2 (LYS+) ura3Δ0 TRP1 leu2Δ0 his3Δ1 ubp9Δ::KanMX* | This study |
| SEN446 | Mata *ADE2 met15Δ0 LYS2 (LYS+) ura3Δ0 TRP1 leu2Δ0 his3Δ1 ubp9Δ::KanMX*<br>*CEN::URA3:: ΔssCPY*-MYC* | This study |
| SEN459 | Mata *ADE2 met15Δ0 LYS2 (LYS+) ura3Δ0 TRP1 leu2Δ0 his3Δ1 ubp14Δ::KanMX*<br>*CEN::URA3* | This study |
| SEN460 | Mata *ADE2 met15Δ0 LYS2 (LYS+) ura3Δ0 TRP1 leu2Δ0 his3Δ1 doa4Δ::KanMX*<br>*CEN::URA3* | This study |
| SEN461 | Mata *ade2-101 met2 lys2-801 ura3-52 trp1::hisG leu2Δ his3Δ200*<br>*CEN::URA3:: ΔssCPY*-MYC* | This study |
| SEN463 | Mata *ade2-101 met2 lys2-801 ura3-52 trp1::hisG leu2Δ his3Δ200 dfm1Δ::KanMX*<br>*CEN::URA3:: ΔssCPY*-MYC* | This study |
| SEN464 | Mata *ADE2 met15Δ0 LYS2 (LYS+) ura3Δ0 TRP1 leu2Δ0 his3Δ1 ubp14Δ::KanMX*<br>*CEN::URA3:: ΔssCPY*-MYC* | This study |
| SEN449 | Mata *ADE2 met15Δ0 LYS2 (LYS+) ura3Δ0 TRP1 leu2Δ0 his3Δ1 ubp6Δ::KanMX*<br>*CEN::URA3* | This study |
| SEN450 | Mata *ADE2 met15Δ0 LYS2 (LYS+) ura3Δ0 TRP1 leu2Δ0 his3Δ1 ubp14Δ::KanMX*<br>*CEN::URA3* | This study |
| SEN451 | Mata *ade2-101 met2 lys2-801 ura3-52 trp1::hisG leu2Δ his3Δ200 dfm1Δ::KanMX*<br>*CEN::URA3* | This study |
| SEN215 | *Mata ADE2::URA3::pTDH3-HMG2-GFPx met15Δ0 LYS2(LYS+) ura3Δ0 TRP1 leu2Δ0 his3Δ1 dfm1Δ::KanMX*<br>*CEN::LEU2::pDFM1-DFM1-3HA-L64V* | Nejatfard, et al., 2021 |

| | | |
|---|---|---|
| SEN216 | *Mata ADE2::URA3::pTDH3-HMG2-GFPx met15Δ0 LYS2(LYS+) ura3Δ0 TRP1 leu2Δ0 his3Δ1 dfm1Δ::KanMX CEN::LEU2::pDFM1-DFM1-3HA-F107S* | Nejatfard, et al., 2021 |
| SEN217 | *Mata ADE2::URA3::pTDH3-HMG2-GFPx met15Δ0 LYS2(LYS+) ura3Δ0 TRP1 leu2Δ0 his3Δ1 dfm1Δ::KanMX CEN::LEU2::pDFM1-DFM1-3HA-K67E* | Nejatfard, et al., 2021 |
| SEN218 | *Mata ADE2::URA3::pTDH3-HMG2-GFPx met15Δ0 LYS2(LYS+) ura3Δ0 TRP1 leu2Δ0 his3Δ1 dfm1Δ::KanMX CEN::LEU2::pDFM1-DFM1-3HA-Q101R* | Nejatfard, et al., 2021 |
| SEN219 | *Mata ADE2::URA3::pTDH3-HMG2-GFPx met15Δ0 LYS2(LYS+) ura3Δ0 TRP1 leu2Δ0 his3Δ1 dfm1Δ::KanMX CEN::LEU2::pDFM1-DFM1-3HA-F58S* | Nejatfard, et al., 2021 |
| SEN529 | *Mata ADE2::URA3::pTDH3-HMG2-GFPx met15Δ0 LYS2(LYS+) ura3Δ0 TRP1 leu2Δ0 his3Δ1 dfm1Δ::KanMX CEN::LEU2* | Nejatfard, et al., 2021 |
| SEN530 | *Mata ADE2::URA3::pTDH3-HMG2-GFPx met15Δ0 LYS2(LYS+) ura3Δ0 TRP1 leu2Δ0 his3Δ1 dfm1Δ::KanMX CEN::LEU2::pDFM1-DFM1-3HA-AR* | Nejatfard, et al., 2021 |
| SEN532 | *Mata ADE2::URA3::pTDH3-HMG2-GFPx met15Δ0 LYS2(LYS+) ura3Δ0 TRP1 leu2Δ0 his3Δ1 dfm1Δ::KanMX CEN::LEU2::pDFM1-DFM1-3HA-AxxxG* | Nejatfard, et al., 2021 |
| SEN534 | *Mata ADE2::URA3::pTDH3-HMG2-GFPx met15Δ0 LYS2(LYS+) ura3Δ0 TRP1 leu2Δ0 his3Δ1 dfm1Δ::KanMX CEN::LEU2::pDFM1-DFM1-3HA* | Nejatfard, et al., 2021 |
| SEN535 | *Mata ADE2::URA3::pTDH3-HMG2-GFPx met15Δ0 LYS2(LYS+) ura3Δ0 TRP1 leu2Δ0 his3Δ1 dfm1Δ::KanMX CEN::LEU2::pDFM1-DFM1-3HA-5aShp* | Nejatfard, et al., 2021 |
| SEN506 | Mata *ade2::ADE2::HIS3 met2 lys2-801 ura3-52, trp1::hisG leu2::LEU2::ADE2:: pADH1-Derlin-1-MYC his3Δ200 dfm1Δ::KanMX* | This study |
| SEN507 | Mata *ade2::ADE2::HIS3 met2 lys2-801 ura3-52, trp1::hisG leu2::LEU2::ADE2:: pADH1-Derlin-1-MYC his3Δ200* | This study |
| SEN510 | Mata *ade2::ADE2::HIS3 met2 lys2-801 ura3-52, trp1::hisG leu2::LEU2::ADE2:: pADH1-Derlin-2-MYC his3Δ200* | This study |
| SEN512 | Mata *ade2::ADE2::HIS3::pGAL1::Hmg2-GFP met2 lys2-801 ura3-52, trp1::hisG leu2::LEU2::ADE2::pADH1-Derlin-2-MYC his3Δ200 dfm1Δ::KanMX* | This study |
| SEN515 | Mata *ade2::ADE2::HIS3::pGAL1::Hmg2-GFP met2 lys2-801 ura3-52, trp1::hisG leu2::LEU2::ADE2 his3Δ200 dfm1Δ::KanMX* | This study |
| SEN516 | Mata *ade2::ADE2::HIS3::pGAL1::Hmg2-GFP met2 lys2-801 ura3-52, trp1::hisG leu2::LEU2::ADE2 his3Δ200* | This study |
| SEN470 | Mata *ade2::ADE2::HIS3 met2 lys2-801 ura3-5, trp1::hisG leu2Δ his3Δ200 CEN::URA3::CFTR-HA* | This study |

| | | |
|---|---|---|
| SEN472 | Mata *ade2::ADE2::HIS3 met2 lys2-801 ura3-52, trp1::hisG leu2::LEU2::ADE2 his3Δ200 dfm1Δ::KanMX CEN::URA3::CFTR-HA* | This study |
| SEN474 | Mata *ade2::ADE2::HIS3 met2 lys2-801 ura3-5, trp1::hisG leu2Δ his3Δ200 CEN::URA3::CFTR-HA-ΔF508* | This study |
| SEN476 | Mata *ade2::ADE2::HIS3 met2 lys2-801 ura3-52, trp1::hisG leu2::LEU2::ADE2 his3Δ200 dfm1Δ::KanMX CEN::URA3::CFTR-HA-ΔF508* | This study |
| SEN478 | Mata *ade2::ADE2::HIS3 met2 lys2-801 ura3-5, trp1::hisG leu2Δ his3Δ200 CEN::URA3::A1PiZ* | This study |
| SEN480 | Mata *ade2::ADE2::HIS3 met2 lys2-801 ura3-52, trp1::hisG leu2::LEU2::ADE2 his3Δ200 dfm1Δ::KanMX CEN::URA3::A1PiZ* | This study |
| SEN452 | Mata *ade2::ADE2::HIS3 met2 lys2-801 ura3-52, trp1::hisG leu2::LEU2::ADE2 his3Δ200 dfm1Δ::KanMX CEN::URA3* | This study |
| SEN455 | Mata *ade2::ADE2::HIS3 met2 lys2-801 ura3-52, trp1::hisG leu2::LEU2::ADE2 his3Δ200 CEN::URA3* | This study |
| SEN365 | ata *ADE2::HIS3::pGAL1::Hmg2-GFP met15Δ0 LYS2 (LYS+) ura3Δ0 TRP1 leu2Δ0 his3Δ1 pdr1Δ::KanMX* | This study |
| SEN366 | Mata *ADE2::HIS3 met15Δ0 LYS2 (LYS+) ura3Δ0 TRP1 leu2Δ0 his3Δ1 pdr1Δ::KanMX* | This study |
| SEN395 | Mata *ADE2::HIS3::pGAL1::Hmg2-GFP met15Δ0 LYS2 (LYS+) ura3Δ0 TRP1 leu2Δ0 his3Δ1 ubp8Δ::KanMX* | This study |
| SEN418 | Mata *ADE2::HIS3 met15Δ0 LYS2 (LYS+) ura3Δ0 TRP1 leu2Δ0 his3Δ1 ubp8Δ::KanMX* | This study |
| SEN396 | Mata *ADE2::HIS3::pGAL1::Hmg2-GFP met15Δ0 LYS2 (LYS+) ura3Δ0 TRP1 leu2Δ0 his3Δ1 miy2Δ::KanMX* | This study |
| SEN419 | Mata *ADE2::HIS3 met15Δ0 LYS2 (LYS+) ura3Δ0 TRP1 leu2Δ0 his3Δ1 miy2Δ::KanMX* | This study |
| SEN397 | Mata *ADE2::HIS3::pGAL1::Hmg2-GFP met15Δ0 LYS2 (LYS+) ura3Δ0 TRP1 leu2Δ0 his3Δ1 otu2Δ::KanMX* | This study |
| SEN420 | Mata *ADE2::HIS3 met15Δ0 LYS2 (LYS+) ura3Δ0 TRP1 leu2Δ0 his3Δ1 otu2Δ::KanMX* | This study |
| SEN398 | Mata *ADE2::HIS3::pGAL1::Hmg2-GFP met15Δ0 LYS2 (LYS+) ura3Δ0 TRP1 leu2Δ0 his3Δ1 ubp2Δ::KanMX* | This study |
| SEN421 | Mata *ADE2::HIS3 met15Δ0 LYS2 (LYS+) ura3Δ0 TRP1 leu2Δ0 his3Δ1 ubp2Δ::KanMX* | This study |
| SEN399 | Mata *ADE2::HIS3::pGAL1::Hmg2-GFP met15Δ0 LYS2 (LYS+) ura3Δ0 TRP1 leu2Δ0 his3Δ1 ubp5Δ::KanMX* | This study |
| SEN422 | Mata *ADE2::HIS3 met15Δ0 LYS2 (LYS+) ura3Δ0 TRP1 leu2Δ0 his3Δ1 ubp5Δ::KanMX* | This study |

| | | |
|---|---|---|
| SEN400 | Mata *ADE2::HIS3::pGAL1::Hmg2-GFP met15Δ0 LYS2 (LYS+) ura3Δ0 TRP1 leu2Δ0 his3Δ1 miy1Δ::KanMX* | This study |
| SEN423 | Mata *ADE2::HIS3 met15Δ0 LYS2 (LYS+) ura3Δ0 TRP1 leu2Δ0 his3Δ1 miy1Δ::KanMX* | This study |
| SEN402 | Mata *ADE2::HIS3::pGAL1::Hmg2-GFP met15Δ0 LYS2 (LYS+) ura3Δ0 TRP1 leu2Δ0 his3Δ1 ubp1Δ::KanMX* | This study |
| SEN425 | Mata *ADE2::HIS3 met15Δ0 LYS2 (LYS+) ura3Δ0 TRP1 leu2Δ0 his3Δ1 ubp1Δ::KanMX* | This study |
| SEN403 | Mata *ADE2::HIS3::pGAL1::Hmg2-GFP met15Δ0 LYS2 (LYS+) ura3Δ0 TRP1 leu2Δ0 his3Δ1 ubp11Δ::KanMX* | This study |
| SEN426 | Mata *ADE2::HIS3 met15Δ0 LYS2 (LYS+) ura3Δ0 TRP1 leu2Δ0 his3Δ1 ubp11Δ::KanMX* | This study |
| SEN405 | Mata *ADE2::HIS3::pGAL1::Hmg2-GFP met15Δ0 LYS2 (LYS+) ura3Δ0 TRP1 leu2Δ0 his3Δ1 ubp7Δ::KanMX* | This study |
| SEN428 | Mata *ADE2::HIS3 met15Δ0 LYS2 (LYS+) ura3Δ0 TRP1 leu2Δ0 his3Δ1 ubp7Δ::KanMX* | This study |
| SEN406 | Mata *ADE2::HIS3::pGAL1::Hmg2-GFP met15Δ0 LYS2 (LYS+) ura3Δ0 TRP1 leu2Δ0 his3Δ1 ubp3Δ::KanMX* | This study |
| SEN429 | Mata *ADE2::HIS3 met15Δ0 LYS2 (LYS+) ura3Δ0 TRP1 leu2Δ0 his3Δ1 ubp3Δ::KanMX* | This study |
| SEN453 | Mata *ade2::ADE2::HIS3 met2 lys2-801 ura3-52, trp1::hisG leu2Δ his3Δ200 CEN::URA3::pCUP1-HBT-Ubiquitin* | This study |
| SEN454 | Mata *ade2::ADE2::HIS3 met2 lys2-801 ura3-52, trp1::hisG leu2Δ his3Δ200 CEN::URA3* | This study |
| SEN457 | Mata *ade2::ADE2::HIS3 met2 lys2-801 ura3-52, trp1::hisG leu2Δ his3Δ200 CEN::URA3::pCUP1-HBT-Ubiquitin* | This study |
| SEN455 | Mata *ade2::ADE2::HIS3 met2 lys2-801 ura3-52, trp1::hisG leu2Δ his3Δ200 CEN::URA3* | This study |
| SEN481 | Mata *ade2::ADE2::HIS3::pGAL1::Hmg2-GFP met2 lys2-801 ura3-52, trp1::hisG leu2Δ his3Δ200 dfm1Δ::KanMX CEN::URA3::pCUP1-HBT-Ubiquitin* | This study |
| SEN456 | Mata *ade2::ADE2::HIS3::pGAL1::Hmg2-GFP met2 lys2-801 ura3-52, trp1::hisG leu2Δ his3Δ200 dfm1Δ::KanMX CEN::URA3* | This study |
| SEN482 | Mata *ade2::ADE2::HIS3 met2 lys2-801 ura3-52, trp1::hisG leu2Δ his3Δ200 dfm1Δ::KanMX CEN::URA3::pCUP1-HBT-Ubiquitin* | This study |
| SEN452 | Mata *ade2::ADE2::HIS3 met2 lys2-801 ura3-52, trp1::hisG leu2Δ his3Δ200 dfm1Δ::KanMX CEN::URA3* | This study |

| | | |
|---|---|---|
| SEN122 | *ade2::ADE2::HIS3 met2 lys2-801 ura3-52::URA3::4xUPRE::GFP TRP1 leu2Δ his3Δ200 der1Δ::CloNat* | This study |
| SEN123 | *ade2::ADE2::HIS3::pGAL::Hmg2-6MYC met2 lys2-801 ura3-52::URA3::4xUPRE::GFP TRP1 leu2Δhis3Δ200 der1Δ::CloNat* | This study |
| RHY11923 | Mata *ade2-101 met2 lys2-801 ura3-5, trp1::hisG::pGAL::Hmg2-GFP leu2Δ his3Δ200 pdr5Δ::HIS3* | This study |
| RHY11924 | Mata *ade2-101 met2 lys2-801 ura3-5, trp1::hisG::pGAL::Hmg2-GFP leu2Δ his3Δ200 pdr5Δ::HIS3 hrd1Δ::KanMX* | This study |
| RHY11925 | Mata *ade2-101 met2 lys2-801 ura3-5, trp1::hisG::pGAL::Hmg2-GFP leu2Δ his3Δ200 pdr5Δ::HIS3 dfm1Δ::CloNAT* | This study |
| RHY11916 | Mata *ade2-101 met2 lys2-801 ura3-5, trp1::hisG::TRP1 leu2Δ his3Δ200 pdr5Δ::HIS3* | This study |
| RHY11917 | Mata *ade2-101 met2 lys2-801 ura3-5, trp1::hisG::TRP1 leu2Δ his3Δ200 pdr5Δ::HIS3 hrd1Δ::KanMX* | This study |
| RHY11918 | Mata *ade2-101 met2 lys2-801 ura3-5, trp1::hisG::TRP1 leu2Δ his3Δ200 pdr5Δ::HIS3 dfm1Δ::CloNAT* | This study |
